# Ethanol’s interaction with BK channel α subunit residue K361 does not mediate behavioral responses to alcohol in mice

**DOI:** 10.1101/2020.10.29.360107

**Authors:** Agbonlahor Okhuarobo, Max Kreifeldt, Pauravi J Gandhi, Catherine Lopez, Briana Martinez, Michal Bajo, Pushpita Bhattacharyya, Alex M Dopico, Marisa Roberto, Amanda J Roberts, Gregg E Homanics, Candice Contet

**Author notes:** Corresponding author Candice Contet Address: The Scripps Research Institute, 10550 N Torrey Pines Road, SR-107, La Jolla, CA 92037, USA.

## Abstract

Large conductance potassium (BK) channels are among the most sensitive molecular targets of ethanol and genetic variations in the channel-forming α subunit have been nominally associated with alcohol use disorders. However, whether the direct action of ethanol at BK α influences the motivation to drink alcohol remains to be determined. In the present study, we sought to investigate the behavioral relevance of this molecular interaction by introducing in the mouse genome a point mutation known to render BK channels insensitive to ethanol while preserving their physiological function. The BK α K361N substitution prevented ethanol from reducing spike threshold in medial habenula neurons. However, it did not alter acute responses to ethanol *in vivo*, including ataxia, sedation, hypothermia, analgesia, and conditioned place preference.

Furthermore, the mutation did not have reproducible effects on alcohol consumption in limited, continuous, or intermittent access home cage two-bottle choice paradigms conducted in both males and females. Notably, in contrast to previous observations made in mice missing BK channel auxiliary β subunits, the BK α K361N substitution had no significant impact on ethanol intake escalation induced by chronic intermittent alcohol vapor inhalation. It also did not affect the metabolic and locomotor consequences of chronic alcohol exposure. Altogether, these data suggest that the direct interaction of ethanol with BK α does not mediate the alcohol-related phenotypes examined here in mice.

## Introduction

Calcium- and voltage-activated, large conductance potassium (BK) channels are one of the primary molecular targets of ethanol in the brain [1–3]. Depending on multiple molecular determinants (e.g., intracellular calcium concentration, alternative splicing, subunit composition, posttranslational modifications, lipid microenvironment), ethanol can either potentiate or reduce BK channel-mediated currents [reviewed in 4]. Polymorphisms in *KCNMA1*, the gene encoding the channel-forming α subunits, were associated with alcohol use disorders (AUD) at nominal significance in several human genomic studies [reviewed in 5]. However, whether the action of ethanol on mammalian BK channels mediates the behavioral effects of ethanol and influences the motivation to drink alcohol remains to be determined. Filling this gap of knowledge has critical implications for the understanding and treatment of AUD.

Until now, the contribution of BK channels to the behavioral effects of ethanol has been studied by genetically manipulating the channel-forming α subunit in worms and flies, and the auxiliary β subunits in mice. Studies in invertebrates showed that BK α mediates the intoxicating effects of ethanol in worms [6, 7] and rapid tolerance to ethanol-induced sedation and increased seizure susceptibility in flies [8–11]. In mice, deletion of BK β4 promotes rapid tolerance to the locomotor depressant effect of ethanol [12] and attenuates ethanol drinking escalation in mice exposed to chronic intermittent ethanol (CIE) vapor inhalation [13]. Conversely, deletion of BK β1 accelerates drinking escalation in CIE-exposed mice [13] and reduces chronic tolerance to ethanol-induced sedation and hypothermia [14]. These findings suggest that BK auxiliary subunits play a role in the adaptive response to chronic ethanol exposure in mammals but fail to provide a direct insight into the role of ethanol’s interaction with BK α subunit in alcohol-related behaviors.

In the present study, we sought to establish whether the action of ethanol at BK α contributes to alcohol drinking and other responses elicited by alcohol in mice. We first used pharmacological modulators of BK channels and then turned to a more specific genetic approach to block the interaction of ethanol with BK α without affecting basal BK channel function. Using recombinant BK channels expressed in *Xenopus* oocytes, residue K361 in BK α cytoplasmic tail domain was shown to play a key role in ethanol recognition as hydrogen bond donor and for the ensuing increase in the channel’s open probability [15]. Importantly, while the K361N substitution confers refractoriness to 100 mM ethanol, it does not alter basal steady-state activity of BK channels, nor their sensitivity to the BK channel primary endogenous activators: voltage and intracellular calcium [15]. It therefore represents a powerful tool to probe the *in vivo* relevance of ethanol’s direct interaction with BK α using gene editing in mice. Our results demonstrate that preventing ethanol from interacting with BK α K361 does not alter several behavioral effects of acute or chronic alcohol exposure, including the motivation to drink alcohol.

## Materials and Methods

### Animals

C57BL/6J mice were obtained from The Jackson Laboratory or from The Scripps Research Institute (TSRI) rodent breeding colony. BK α K361N knockin (KI) mice were generated at the University of Pittsburgh. Breeders were sent to TSRI, where a colony was maintained by mating heterozygous (Het) males and females such that experimental mice were littermates. KI mice were backcrossed to C57BL/6J mice every 1-2 years to prevent genetic drift.

Mice were maintained on a 12 h/12 h light/dark cycle. Food (Teklad LM-485, Envigo) and acidified or reverse osmosis purified water were available *ad libitum*. Sani-Chips (Envigo) were used for bedding substrate. Behavioral testing was started when mice were at least 10 weeks old. Mice were individually housed for drinking experiments and metabolic and activity tracking, but group-housed otherwise. Testing was conducted during the dark phase of the circadian cycle, except for conditioned place preference, which was conducted during the light phase.

All procedures adhered to the National Institutes of Health Guide for the Care and Use of Laboratory Animals and were approved by the Institutional Animal Care and Use Committees of the University of Pittsburgh and TSRI.

### Experimental cohorts

Independent cohorts of C57BL/6J males were used to test the effects of penitrem A on tremors (n=28, 6-8 mice per dose), alcohol drinking (n=20) and saccharin drinking (n=20), as well as the effects of paxilline (n=18) and BMS-204352 (n=30) on alcohol drinking.

To evaluate the effect of BK α K361N substitution, body weights were collected in 6-week-old males (WT, n=7; Het, n=12; KI, n=6). Electrophysiological recordings were obtained from experimentally naïve adult males (WT paxilline, n=6 cells from 3 mice; WT ethanol, n=11 cells from 4 mice; KI ethanol, n=9 cells from 4 mice). Separate cohorts of males were tested for ethanol clearance rate (WT, n=3; Het, n=4; KI, n=4), ethanol-induced ataxia, sedation, and hypothermia (WT, n=10; Het, n=11; KI, n=8), ethanol-induced analgesia (WT, n=7; Het, n=9; KI, n=7), and conditioned place preference (WT, n=11; KI, n=15). Three independent cohorts of mice, each containing an equivalent number of WT and KI males, were tested for limited-access alcohol drinking acquisition and CIE-induced escalation, and their data were pooled for analysis (WT, n=19; KI, n=15). This experiment was repeated in another set of three independent cohorts of mice, each containing an equivalent number of WT and KI males and females, and the data were pooled by sex for analysis (WT males, n=21; KI males, n=20; WT females, n=21; KI females, n=22). Two additional cohorts of males were used to measure metabolism/activity (WT, n=8; KI, n=7) and circadian rhythmicity (WT, n=8; KI, n=8) during withdrawal from CIE. Continuous- and intermittent-access alcohol drinking were first tested in a cohort containing both males and females (WT males, n=5; KI males, n=7; WT females, n=6; KI females, n=6), and later repeated in a separate cohort of females (WT females, n=10; KI females, n=9).

### Drugs

Penitrem A was purchased from Sigma (P3053) for tremor assessment and from Enzo Life Sciences (BML-KC157) for drinking experiments. It was dissolved in dimethylsulfoxide (DMSO) at 10 mg/mL and diluted in saline for intraperitoneal (i.p.) injection (0.1 mL per 10 g body weight). The final concentration of DMSO was 50% for tremor assessment and 10% for drinking experiments.

Paxilline was purchased from Sigma (P2928), dissolved in DMSO at 10 mM and diluted in phosphate-buffered saline (137 mM NaCl, 2.7 mM KCl, 1.8 mM KH_2_PO_4_, 10.1 mM Na_2_HPO_4_, pH 7.4) for i.p. injection (1:2000 for 22 µg/kg dose, 1:400 for 110 µg/kg dose, 1:80 for 550 µg/kg dose). Each dose was tested on a different week. Doses were tested in ascending order, and vehicle and drug treatments were counterbalanced over two consecutive days for each dose.

This dose range was selected based on pilot testing that indicated reduced mobility at 1.1 mg/kg and tremors at 4.4 mg/kg, which would have confounded drinking behavior, as well as on reported anticonvulsant properties of ultra-low-dose paxilline [16].

BMS-204352 was purchased from Sigma (SML1313), dissolved in DMSO at 16 mg/mL and diluted in Tween-80:saline at a 1:1:80 ratio for i.p. injection. A dose of 2 mg/kg was selected based on its ability to reverse behavioral abnormalities in *Fmr1* mutant mice [17, 18]. This dose did not produce a noticeable effect on mouse behavior (e.g., hyperactivity or sedation).

Ethyl alcohol 200 proof (PHARMCO-AAPER, 111000200) was used for i.p. administration (diluted in saline) and drinking solutions (diluted in acidified or reverse osmosis purified water). Ethyl alcohol 95% (PHARMCO-AAPER, 111000190) was used for vapor inhalation.

### Generation of BK ***α*** K361N KI mice

KI mice were produced using CRISPR/Cas9 technology as previously described in detail [19]. Briefly, a sgRNA targeting *Kcnma1* in exon 9 near the intended mutation site was identified using the CRISPR Design Tool [20]. Two overlapping PCR primers (forward: GAAATTAATACG ACTCACTATAGGAGTGTCTCTAACTTCCTGAGTTTTAGAGCTAGAAATAGC; R: AAAAGCA CCGACTCGGTGCCACTTTTTCAAGTTGATAACGGACTAGCCTTATTTTAACTTGCTATTTCT AGCTCTAAAAC) were used to generate a T7 promoter containing sgRNA template as described [21]. The sgRNA and Cas9 mRNA were produced by *in vitro* translation, purified (MEGAclear Kit, Ambion), ethanol precipitated, and resuspended in DEPC treated water. A 120- nucleotide single stranded DNA repair template oligonucleotide harboring the desired mutations in exon 9 of *Kcnma1* was purchased as Ultramer DNA (Integrated DNA Technologies, Coralville, IA). sgRNA (25 ng/µl), Cas9 mRNA (50 ng/µl), and repair oligo (100 ng/µl) were combined and injected into the cytoplasm of C57BL/6J one-cell embryos as described [22].

Embryos that survived injection were transferred to the oviduct of day 0.5 post-coitum pseudo- pregnant CD-1 recipient females. Pups resulting from injected embryos were screened for DNA sequence changes in exon 9 of the *Kcnma1* gene by PCR/DNA sequence analysis. A male founder mouse harboring the desired changes was mated to C57BL/6J females to establish the KI line. The *Kcnma1* exon 9 containing amplicon from all Het F1 animals that were shipped for the TSRI breeding colony were sequenced to verify the fidelity of the mutated locus. The founder mouse harbored no off-target mutations (data not shown) in any of the top 7 off-target sites predicted by the Off –Targets tool of the Cas9 Online Designer [23].

Mice were genotyped by subjecting tail clip lysates to polymerase chain reaction (PCR) using a pair of primers (forward: GCTTTGCCTCATGACCCTCT; reverse: TGAACAAGGGTGCTGCTTC A) that amplifies a 450-bp fragment of the *Kcnma1* gene. The PCR products were then digested with Tru1I and the resulting fragments were visualized by electrophoresis in an ethidium bromide-stained agarose gel. Tru1I digestion yields two fragments (107 + 343 bp) in the wild- type (WT) allele and three fragments (107 + 149 + 194 bp) in the KI allele (see KI-specific Tru1I site in **Fig. 2A**).

To verify that the mutation was also present in *Kcnma1* mRNA, RNA was isolated from a KI mouse brain hemisphere using the RNeasy Plus Universal Mini Kit (Qiagen, 73404), 2 µg of RNA was reverse-transcribed using the Transcriptor First Strand cDNA Synthesis Kit with random hexamer primers (Roche, 04379012001), and a 370-bp fragment (nucleotides 1304- 1673 of NM_010610.3) was amplified from the resulting cDNA. This fragment was cloned into pBluescript II and sequenced with a T3 primer (Genewiz).

### In situ hybridization

Custom locked nucleic acid (LNA) oligoprobes labeled with digoxigenin in 3’ and 5’ were obtained from Exiqon (Denmark). The sequences were as follows:

STREX (batch 620646) /5DigN/ATGCTCGTCTCATTCTCTTGTA/3Dig_N/ ALCOREX (batch 620645) /5DigN/TTCCTGATTGACTGATACCA/3Dig_N/

The same protocol and reagents as in [24] were used to prepare brain sections and label *Kcnma1* splice variants. The probes were hybridized overnight at 48°C at a final concentration of 5 nM.

### Electrophysiological recordings

Recordings were obtained from sagittal brain slices (300-μm thick) containing the medial habenula (mHb). The slices were cut in oxygenated (95% O_2_/5% CO_2_), ice-cold high-sucrose solution (pH 7.3-7.4, 206.0 mM sucrose, 2.5 mM KCl, 0.5 mM CaCl_2_, 7.0 mM MgCl_2_, 1.2 mM NaH_2_PO_4_, 26 mM NaHCO_3_, 5 mM glucose, 5 mM HEPES) using a vibratome (Leica VT1200S) and then incubated in an oxygenated recovery solution (130 mM NaCl, 3.5 mM KCl, 2 mM CaCl_2_, 1.25 mM NaH_2_PO_4_, 1.5 mM MgSO_4_, 24 mM NaHCO_3_, 10 mM glucose, and 1 mM kynurenic acid) at 32°C for 30 min. After recovery, the slices were kept in oxygenated artificial cerebrospinal fluid (aCSF: 130 mM NaCl, 3.5 mM KCl, 2 mM CaCl_2_, 1.25 mM NaH_2_PO_4_, 1.5 mM MgSO_4_, 24 mM NaHCO_3_, and 10 mM glucose) at room temperature for at least 30 min until recording started.

mHb neurons were visualized using an upright microscope (Olympus BX51WI) with infrared-differential interference contrast (IR-DIC) optics, a w40 or w60 water immersion objective, and a CCD camera (EXi Aqua, QImaging). Whole-cell recordings of the current-voltage relationships (IVs) were performed in the current clamp mode at a sampling rate per signal of 50 kHz and low-pass filtered at 10 kHz, using a Multiclamp 700B amplifier, Digidata 1440A and pClamp 10.2 software (all Molecular Devices). The neurons were held at -60 mV, and the recording glass electrodes (3-6 MΩ resistance) were filled with K-gluconate internal solution: 145 mM K- gluconate, 5 mM EGTA, 2 mM MgCl2, 10 mM HEPES, 2 mM Mg+-ATP, 0.2 mM Na+-GTP. The protocol consisted of a set of 15 depolarizing steps lasting 400 ms (Δ +10 pA, starting at -20 pA). The recordings were conducted before (baseline) and during a drug application. The BK channel blocker paxilline (300 nM) and ethanol (50 mM) were bath applied for 8-12 min and 6-8 min, respectively. The recorded IVs were analyzed by NeuroExpress 19.0 [25].

### Tremors

Tremors were scored according to the following scale [26]: 0 = no tremor; 1 = no resting tremor, short-lasting low-intensity shaking elicited by handling; 2 = no resting tremor, continuous low-intensity shaking elicited by handling; 3 = spontaneous low-intensity tremor, aggravated by handling; 4 = severe spontaneous tremor, convulsive episode elicited by handling; score 5 was not observed.

### Ethanol clearance rate

Mice were i.p. injected with 2 g/kg ethanol (20% v:v, 0.13 ml/10 g body weight). Tail vein blood was collected 30 min, 90 min and 180 min later and processed for blood ethanol concentration (BEC) determination by gas chromatography and flame ionization detection (Agilent 7820A).

### Motor coordination and ethanol-induced ataxia

Motor coordination was evaluated using an AccuRotor rotarod (Accuscan Instruments) accelerating from 4 to 40 rpm over 300 s. Mice were positioned on the rotating rod and speed at fall (rpm) was recorded. For motor learning, mice were subjected to 5 trials per day (30-90 min apart) for 3 consecutive days. For ataxia testing, the rod was rotating at a constant speed of 8 rpm and the mice had to stay on the rod for at least 30 s to pass. Ataxia testing was conducted 4-5 days after the last training trial and all mice were able to pass the criterion. They were then i.p. injected with 1.5 g/kg ethanol (0.1 mL/10 g body weight) and tested approximately every 4 min until they were able to pass the criterion again. At this point, blood was collected from the retroorbital sinus and processed for BEC determination using a GM7 analyzer (Analox Instruments).

### Ethanol-induced sedation and hypothermia

Baseline body temperatures were first determined using a MicroTherma 2K thermometer (ThermoWorks) fitted with a rectal probe. Mice were then i.p. injected with 3.5 g/kg ethanol (0.2 mL/10 g body weight), which resulted in loss of righting response (i.e., sedation). Mice were placed on their backs and the time at which each mouse regained its righting response was recorded. At this point, retroorbital blood was sampled and BECs were determined using a GM7 analyzer. Body temperatures were again recorded 60 and 120 min after injection.

### Ethanol-induced analgesia

A digital Randall-Selitto apparatus (Harvard Apparatus 76-0234) was used to measure mechanical nociceptive thresholds, as described in [27]. The mouse was habituated to enter a restrainer made of woven wire (stainless steel 304L 200 mesh, Shanghai YiKai) over the course of 3 days. On testing days, the mouse was gently introduced into the restrainer and the distal portion of the tail was positioned under the conic tip of the apparatus. The foot switch was then depressed to apply uniformly increasing pressure onto the tail until the first nociceptive response (struggling or squeaking) occurred. The force (in g) eliciting the nociceptive response was recorded. A cutoff force of 600 g was enforced to prevent tissue damage. The measure was repeated on the medial and proximal parts of the tail of the same mouse, with at least 30 seconds between each measure. The average of the three measures (distal, medial, proximal) was used as nociceptive value for that day. The analgesic effect of ethanol was tested over 4 consecutive days using a Latin square design. Testing was conducted 5 min after i.p. injection of 20% v:v ethanol (0, 1.5, 2 and 2.5 g/kg, 0.1-0.17 mL/10 g body weight).

### Ethanol conditioned place preference

The apparatus was made of matte black acrylic and consisted of a 42 cm long x 21 cm wide x 31 cm high rectangular box (inner dimensions) with a removable central divider (ePlastics, San Diego). In one compartment, the floor was covered with coarse mesh (stainless steel 304L 10 mesh, Shanghai YiKai) and the walls were decorated with white discs (5-cm dot sticker, ChromaLabel). In the other compartment, the floor was smooth and the walls were uniformly black. Pre-conditioning, conditioning, and post-conditioning trials were conducted on consecutive days, 2 h into the light phase of the circadian cycle. During the pre-conditioning and post-conditioning tests, mice had access to both compartments during 15 min and their motion was video-recorded by a ceiling-mounted camera connected to ANY-maze (Stoelting Co.).

During the conditioning trials, the mice were i.p. injected with saline or 2 g/kg ethanol (20% v:v, 0.13 mL/10 g body weight) and immediately confined to the compartment paired with this treatment during 30 min. A biased design was used to assign compartments to saline or ethanol for each mouse, i.e., ethanol was always assigned to the least favorite compartment (mesh floor for 6 WT and 9 KI mice, smooth floor for 5 WT and 6 KI mice). Treatments were alternated for a total of 8 conditioning trials (4 saline and 4 ethanol) and the order of treatment was counterbalanced within each genotype. Conditioned place preference was reflected by an increase in the time spent in the ethanol-paired compartment after *vs*. before conditioning.

### Alcohol drinking

Mice were single-housed 3 days before testing started and remained single-housed throughout the duration of all drinking experiments. Voluntary alcohol consumption was assessed as a two-bottle choice (2BC) between water and ethanol in the home cage. Ethanol concentration was 15% v:v as in [28] and [29], except for the continuous/intermittent access repeat experiment, which used 20% w:v as in [30]. For limited access, 2-h 2BC sessions were started at the beginning of the dark phase (except for the penitrem A study, in which sessions were started 2 h into the dark phase) and conducted Mon-Fri. For continuous access, bottles were weighed daily 3 h into the dark phase. For intermittent access, the ethanol bottle was present for 24-h periods starting 3 h into the dark phase on Mon, Wed, and Fri, and replaced with a second water bottle the rest of the time. In all paradigms, the positions of the ethanol and water bottles were alternated each day to control for side preference. Ethanol intake was determined by weighing bottles before and after the session, subtracting the weight lost in bottles placed in an empty cage (to control for spill/evaporation) and dividing by the mouse bodyweight (measured weekly). A similar procedure was used to assess 2-h saccharin (0.005% w:v) consumption in the penitrem A study. Ethanol intake is expressed as g of ethanol molecule, while saccharin intake is expressed as g of saccharin solution; both are normalized to the mouse’s body weight. CIE vapor inhalation was used to increase voluntary ethanol intake in 2-h 2BC sessions, as described in [13, 28]. Mice were first subjected to two weeks of 2BC (Mon-Fri) and split into two groups of equivalent baseline ethanol intake (average of last three days). Weeks of CIE (or air) inhalation (4 x 16-h intoxication/8-h withdrawal, Mon-Fri) were then alternated with weeks of 2BC (Mon-Fri) for a total of 3-5 rounds.

BK channel modulators were injected 30 min prior to 2-h 2BC.

### Chronic intermittent ethanol (CIE) vapor inhalation

The inhalation chambers were made of sealed plastic mouse cages (Allentown). An electronic metering pump (Iwaki EZB11D1-PC) dripped 95% ethanol into a flask placed on a warming tray at a temperature of 50°C. Drip rate was adjusted to achieve target BECs of 150-250 mg/dL. An air pump (Hakko HK-80L) conveyed vaporized ethanol from the flask to each individual chamber. The air flow was set at a rate of 15 L/min for each pair of chambers. Each chamber was diagonally divided by a mesh partition to provide single housing for two mice. Mice were injected i.p. with ethanol (1.5 g/kg) and pyrazole (68 mg/kg, Sigma-Aldrich, P56607) diluted in saline, in a volume of 0.1 mL/10 g body weight, before each 16-h ethanol vapor inhalation session. Blood was sampled from the caudal vein at the end of a 16-h intoxication session. The tip of the tail was nicked with a scalpel blade, blood was collected with a heparinized capillary tube and centrifuged at 13,000 g for 10 min. BECs were measured using a GM7 analyzer or by gas chromatography and flame ionization detection. On CIE weeks, control (Air) mice received pyrazole only. Average BECs in each CIE-exposed cohort are reported in **Table 1**.

**Table 1.**
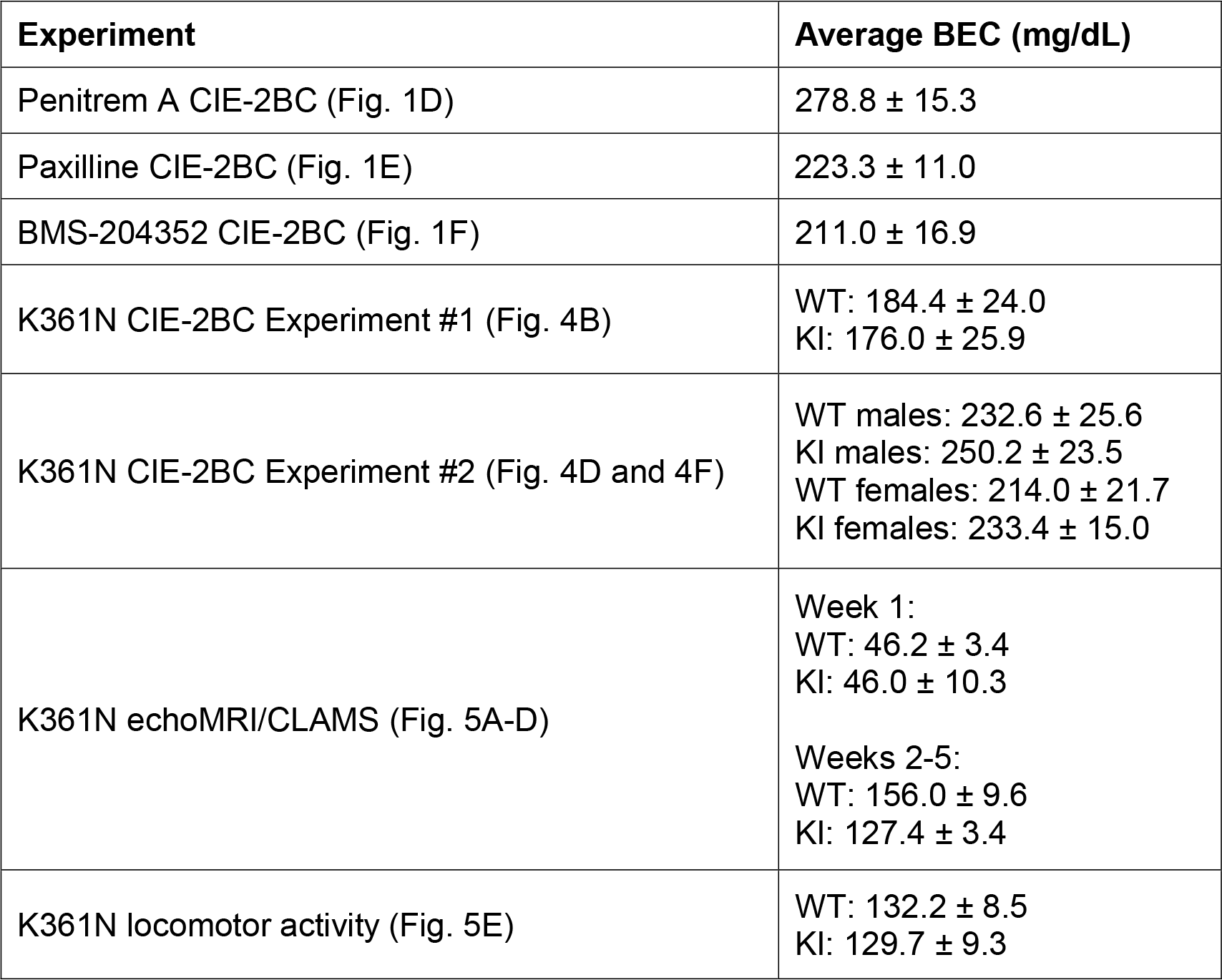
Average blood ethanol concentrations (BEC) during chronic intermittent ethanol (CIE) vapor inhalation.

| Experiment | Average BEC (mg/dL) |
| --- | --- |
| Penitrem A CIE-2BC (Fig. 1D) | 278.8 ± 15.3 |
| Paxilline CIE-2BC (Fig. 1E) | 223.3 ± 11.0 |
| BMS-204352 CIE-2BC (Fig. 1F) | 211.0 ± 16.9 |
| K361N CIE-2BC Experiment #1 (Fig. 4B) | WT: 184.4 ± 24.0<br>KI: 176.0 ± 25.9 |
| K361N CIE-2BC Experiment #2 (Fig. 4D and 4F) | WT males: 232.6 ± 25.6<br>KI males: 250.2 ± 23.5<br>WT females: 214.0 ± 21.7<br>KI females: 233.4 ± 15.0 |
| K361N echoMRI/CLAMS (Fig. 5A-D) | Week 1:<br>WT: 46.2 ± 3.4<br>KI: 46.0 ± 10.3<br>Weeks 2-5:<br>WT: 156.0 ± 9.6<br>KI: 127.4 ± 3.4 |
| K361N locomotor activity (Fig. 5E) | WT: 132.2 ± 8.5<br>KI: 129.7 ± 9.3 |

### Metabolism and actimetry

Mice were exposed to CIE every other week, starting with a priming week at sub-intoxicating BECs, and followed by 4 weeks at intoxicating BECs. Body composition was analyzed by quantitative nuclear magnetic resonance (EchoMRI 3-in-1, EchoMRI LLC) 72 h after the last vapor exposure. Mice were then immediately placed in individual metabolic cages (Comprehensive Laboratory Animal Monitoring System [CLAMS], Oxymax, Columbus Instruments), at the beginning of the dark phase. The following data were collected every 18 min for a total of 108 h: oxygen consumption (VO_2_), carbon dioxide production (VCO_2_), food intake, water intake, and locomotor activity. The respiratory exchange ratio (RER), calculated as VCO_2_/VO_2_, provides an indicator of the substrate being metabolized, ranging from 0.7 when the predominant fuel source is fat to 1 when the predominant fuel source is carbohydrate [31].

Locomotor activity counts (beam interruptions) were used by CLAMS-HC Sleep Detection function to track sleeping bouts, as defined by 4 (or more) consecutive 10-sec epochs with 0 activity counts [32]. The first 12 hours (dark phase) were considered habituation and excluded from analysis. The following 96 h were binned by 12-h light and dark phases and averaged across the 4 days for statistical analysis.

### Circadian rhythmicity

Mice were exposed to CIE every other week for a total of 4 weeks and transferred to individual locomotor activity cages (Photobeam Activity System-Home Cage, San Diego Instruments) 72 h after the last vapor exposure. Mice were maintained on a 12 h/12 h light/dark cycle for 7 consecutive days, then switched to constant darkness for an additional 11 days. Ambulation counts represent consecutive beam breaks (8 x 4 beams in the 18.5” x 10” frame) and were collected in 1-h bins. Chi-square periodogram analysis was conducted in R (‘zeitgebr’ package, https://github.com/rethomics/zeitgebr) to determine the circadian period length and relative power during constant darkness [33, 34], using the last 240 hours of recording and a 6-min resampling rate.

### Statistical analysis

Data were analyzed in Prism 9.5.1 (GraphPad) and Statistica 13.3 (Tibco Software Inc.). Electrophysiological data were analyzed by multiple paired t-tests using a false discovery rate of 1%. Tremor scores were analyzed by Kruskal-Wallis analysis of variance (ANOVA) of the area under the curve, followed by Dunn’s comparisons to vehicle. Alcohol and saccharin drinking was analyzed by repeated-measures (RM) ANOVA with dose (pharmacological experiments), week (2-h 2BC acquisition and escalation, weekly averages), or access schedule (continuous/intermittent 2BC, last 24-h period of each phase) as within-subject variable.

*Posthoc* tests were conducted using Dunnett’s test for comparisons to vehicle, and Šidák’s test otherwise. For escalation, significant week x vapor interactions were followed up with two-way ANOVAs (genotype, vapor) on individual weeks. The effect of genotype on body weight, ataxia, and sedation was analyzed by one-way ANOVA. Ethanol’s clearance rate, hypothermia, analgesia, and place preference were analyzed by two-way RM-ANOVA, with time, dose, or conditioning as within-subject variable and genotype as between-subject variable. EchoMRI data, circadian period length and relative power were analyzed by two-way ANOVA (genotype, vapor). CLAMS data were analyzed by three-way RM-ANOVA, with phase as within-subject variable and genotype and vapor as between-subject variables. When there was a significant interaction between phase and vapor, two-way ANOVAs were further conducted for each phase. The Greenhouse-Geisser correction was used to adjust for lack of sphericity in RM- ANOVAs with three or more levels. Data are expressed as mean ± s.e.m.

## Results

### Non-tremorgenic pharmacological inhibition of BK channel activity can modulate moderate alcohol drinking

We first sought to examine the contribution of BK channels to voluntary ethanol consumption using a pharmacological approach in C57BL/6J males. Since ethanol can activate some neuronal BK channels [1-3, 12, 35], we hypothesized that blocking BK channels may interfere with the motivational properties of ethanol and increase (to overcome BK channel blockade) or decrease (if blockade is unsurmountable) alcohol drinking. On the other hand, in central amygdala GABAergic synapses, ethanol can mimic the effect of BK blockers [36], such that BK channel inhibitors may prime alcohol seeking at low doses and substitute for ethanol (i.e., reduce the motivation to consume alcohol) at higher doses.

We first used penitrem A, a brain-penetrant fungal alkaloid that potently inhibits BK channels [37, 38]. Penitrem A induced tremors in a dose-dependent manner (**Fig. 1A**, main effect of dose: H_3,24_=23.4, p<0.0001; *posthoc* comparisons to vehicle: 0.2 mg/kg, p=0.0055; 0.5 mg/kg, p<0.0001), as reported previously [39]. Consistent with its tremorgenic effect, the dose of 0.2 mg/kg abolished both ethanol (**Fig. 1B**, dose effect: F_1.899,36.08_=65.5, p<0.0001; vehicle *vs.* 0.2 mg/kg, p<0.0001) and saccharin (**Fig. 1C**, dose effect: F_1.864,35.41_=59.6, p<0.0001; vehicle *vs.* 0.2 mg/kg, p<0.0001) drinking. The dose of 0.1 mg/kg reduced ethanol intake (p=0.0169) without affecting saccharin intake (p=0.98). The lowest dose of 0.05 m/kg did not affect ethanol (p=0.57) or saccharin (p=0.98) intake (**Fig. 1B-C**).

**Figure 1.**
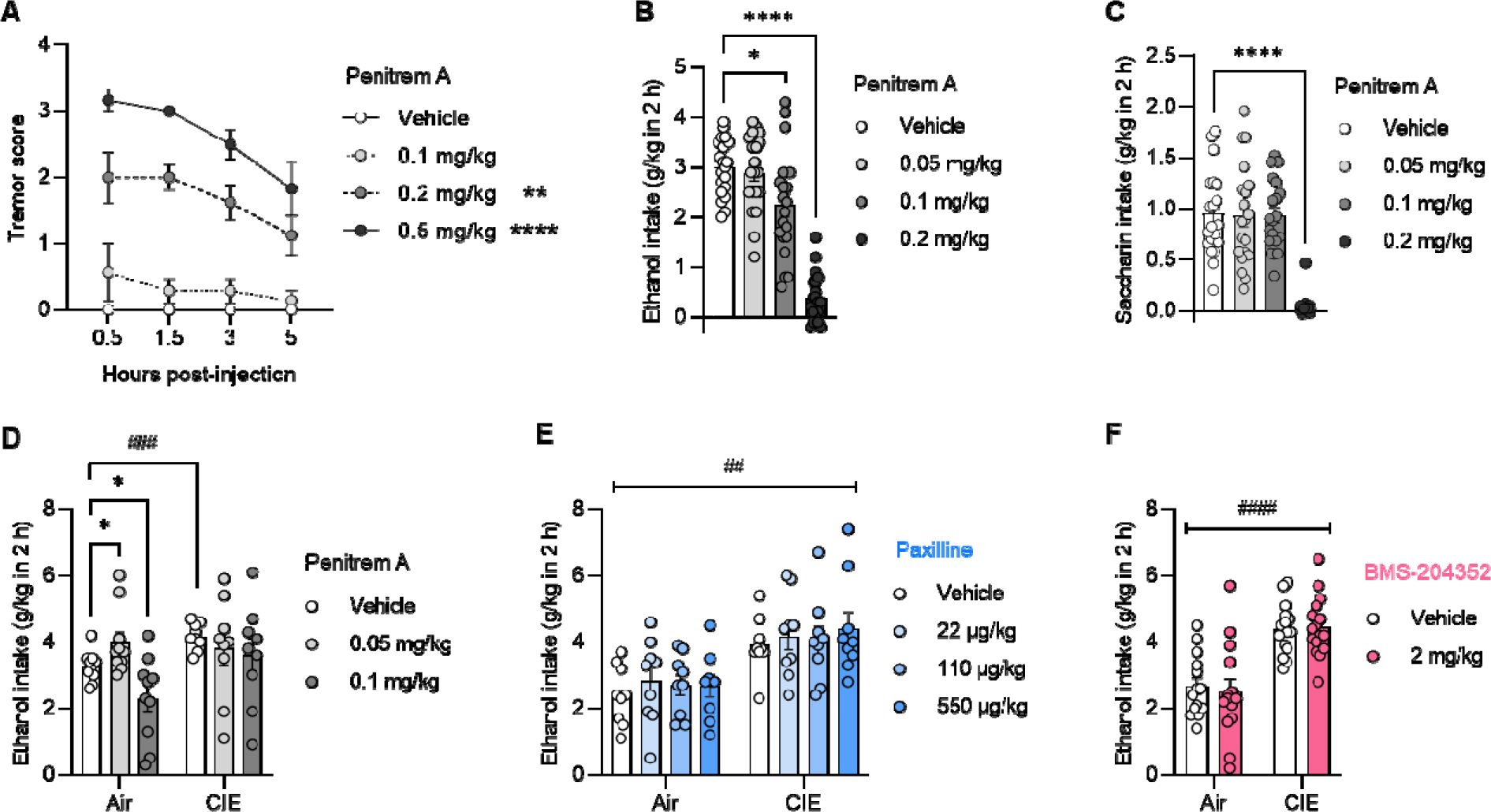
Effects of BK channel pharmacological modulators on baseline and escalated alcohol drinking. A. Penitrem A, a BK channel blocker, induced tremors in a dose-dependent manner in alcohol-naïve C57BL/6J males (between-subject design). Error bars show s.e.m. **B-F.** C57BL/6J male mice were given access to voluntary alcohol (**B, D-F**) or saccharin (**C**) consumption in 2-h two-bottle choice (2BC) sessions. The BK channel blockers penitrem A (**B-D**) or paxilline (**E**) or the BK channel opener BMS-204352 (**F**) were injected i.p. 30 min before 2BC (within-subject design). Some mice were exposed to chronic intermittent ethanol (CIE) vapor inhalation to increase voluntary ethanol intake, compared to mice inhaling air only (Air, between-subject design). Significant difference with vehicle: *, p<0.05; **; ****, p<0.0001. Significant effect of CIE: ^##^, p<0.01; ^###^, p<0.001; ^####^, p<0.0001.

Based on our previous findings in BK β1 and β4 knockout (KO) mice [13], we reasoned that the effect of BK channel modulation may be sensitive to CIE exposure. There was a significant interaction between vapor and penitrem A (F_2,34_=3.47, p=0.0426) whereby vehicle-injected CIE mice consumed more alcohol than their Air-exposed counterparts (p=0.0008) and penitrem A had dose-dependent effects in Air mice (increase at 0.05 mg/kg, p=0.0165; decrease at 0.1 mg/kg, p=0.0355) but not in CIE mice (**Fig. 1D**).

Tremorgenic mycotoxins can inhibit BK channels via different mechanisms and may therefore have a differential effect on ethanol-induced potentiation of BK-mediated currents. Notably, the association of β1 subunits reduces BK channel sensitivity to penitrem A by 10-fold, while it does not affect sensitivity to paxilline, a highly selective BK channel blocker [38, 40]. Since β1 subunits influence ethanol intake escalation in CIE-exposed mice [13], we next tested the effect of paxilline in both Air and CIE mice. We limited our analysis to non-tremorgenic doses (see *Methods* for dose range determination). Paxilline did not affect ethanol intake regardless of vapor exposure (**Fig. 1E**, dose effect: F_1.865,29.84_=1.0, p=0.38; vapor effect: F_1,16_=11.2, p=0.004; dose x vapor interaction: F_3,48_=0.27, p=0.85).

To further investigate the ability of BK channels to modulate ethanol intake, we tested the effect of a BK channel opener, BMS-204352. At 2 mg/kg, a dose that rescues several behavioral deficits of *Fmr1* KO mice [17, 18], BMS-204532 did not impact moderate (Air) or excessive (CIE) ethanol drinking (**Fig. 1E**, treatment effect: F_1,28_=0.09, p=0.77; vapor effect: F_1,28_=28.3, p<0.0001; treatment x vapor interaction: F_1,28_=0.6, p=0.43).

### Generation and validation of BK ***α*** K361N knockin mice

The significance of pharmacological manipulations is inherently limited because they perturb the physiological activity of BK channels rather than selectively targeting their ethanol-sensing capacity. We therefore turned to a genetic approach to probe the role of ethanol’s direct action at BK channels in the motivation to consume alcohol. An asparagine substitution of residue K361 of the mouse BK α subunit was shown to abolish ethanol’s ability to increase BK channel steady-state activity without affecting unitary conductance, calcium sensitivity, or voltage sensitivity, thereby providing a unique opportunity to selectively disrupt the direct effect of ethanol on BK channels [15].

Accordingly, we generated a knockin (KI) mouse expressing the K361N mutant instead of the wildtype (WT) BK α on a C57BL/6J background. A CRISPR/Cas9 strategy was used to introduce two nucleotide mutations in the *Kcnma1* gene: A G-to-T missense mutation modifying the triplet encoding K361 into an asparagine-coding triplet, and a silent G-to-T mutation introducing a Tru1I restriction site to facilitate mouse genotyping (**Fig. 2A**). KI mice were viable, and all three genotypes (KI, Het, and WT) were obtained in Mendelian proportions. The presence of the mutations in the *Kcnma1* mRNA was confirmed by mouse brain cDNA sequencing (**Fig. 2B**). We verified that the basal function of BK channels was preserved in KI mice, based on the known phenotype of mice missing BK α. Accordingly, while BK α KO mice displayed 15-20% smaller body weights than their WT counterparts at 4 and 8 weeks of age [41], there was no effect of the K/N361 genotype on body weight at 6 weeks of age (**Fig. 2C**, F_2,22_=0.4, p=0.70). Furthermore, while BK α KO mice displayed major motor coordination deficits [41], BK α K361N KI mice acquired the accelerating rotarod task at the same rate as their Het and WT counterparts (**Fig. 2D**, effect of trial: F_14,336_=37.2, p<0.0001; effect of genotype: F_2,24_=0.8, p=0.48; trial x genotype interaction: F_28,336_=0.8, p=0.73).

**Figure 2.**
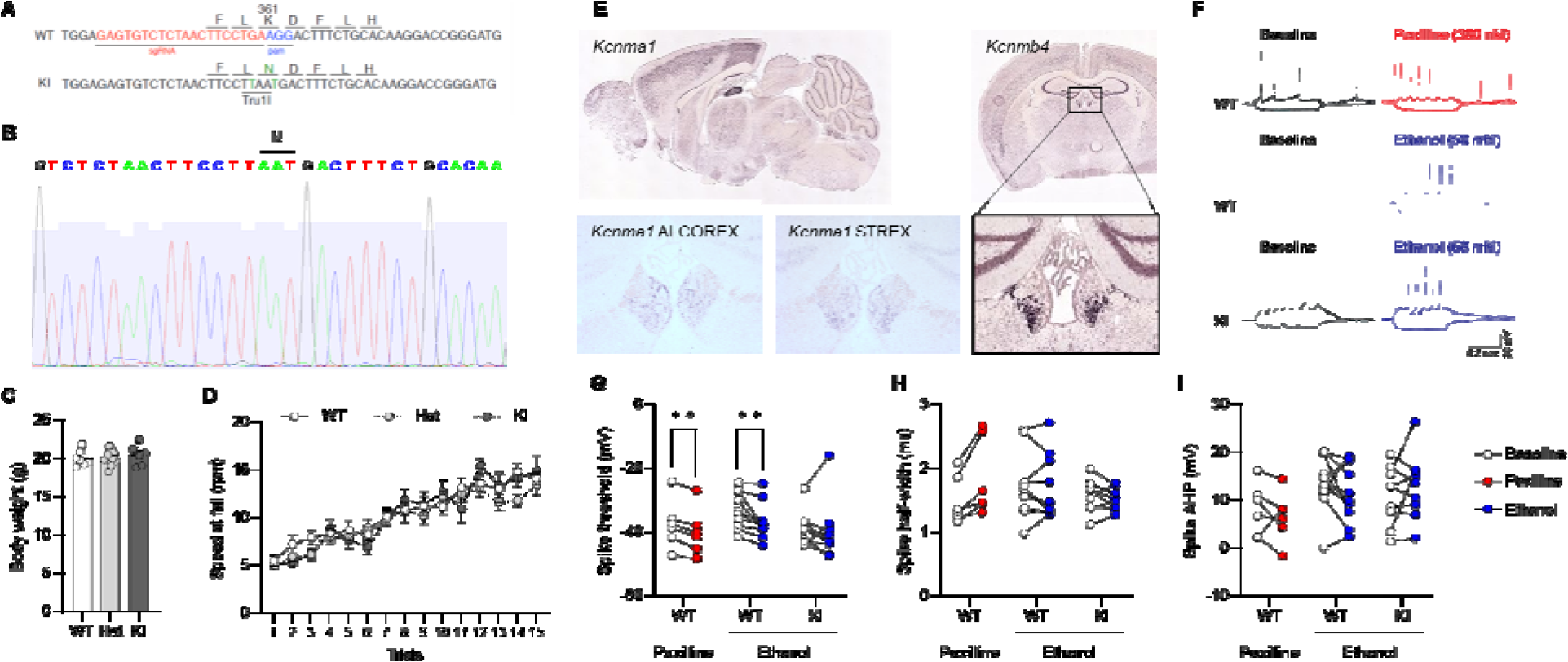
Generation and validation of BK. α **K361N knockin mice. A.** Design of the CRISPR/Cas9 construct used to introduce the K361N substitution in C57BL/6J mice. The single guide RNA (sgRNA) sequence is shown in red, the protospacer adjacent motif (PAM) is shown in blue, and the two mutated nucleotides are shown in green. WT, wildtype allele; KI, knockin allele. **B.** Verification of the mutated sequence in cDNA prepared from the brain of a K361N KI mouse. The triplet encoding the K361N substitution is highlighted. **C.** Body weights measured in males at 6 weeks of age. **D.** Motor coordination measured in the accelerating rotarod assay in adult males. Error bars show s.e.m. There was no effect of genotype on either measure. **E.** Distribution of transcripts encoding BK channel α (*Kcnma1*) and β4 (*Kcnmb4*) (Allen Mouse Brain Atlas, https://mouse.brain-map.org, experiments 74578206 and 72283793), as well as alternatively spliced *Kcnma1* exons ALCOREX and STREX, in mouse brain sections, showing their enrichment in the medial habenula (mHb). **F.** Representative traces from electrophysiological recordings of mHb neurons from WT and KI brain slices before (open circles in **G-I**) and during (closed circles in **G-I**) treatment with paxilline (300 nM, red) or ethanol (50 mM, blue). **G.** Paxilline and ethanol both reduced the spike threshold in WT neurons, but KI neurons were insensitive to ethanol. There was no significant effect of paxilline or ethanol on spike half-width (**H**) or spike afterhyperpolarization (**I**). Baseline vs. drug: **, q<0.01 (FDR 1%).

We next sought to confirm that the K361N substitution imparts ethanol resistance to native BK channels, as seen with recombinant channels expressed in *Xenopus* oocytes.

Electrophysiological recordings were obtained from the medial habenula (mHb), a brain region that expresses high levels of BK channel subunits (*Kcnma1* and *Kcnmb4* transcripts), including an alternatively spliced *Kcnma1* exon known to confer sensitivity to ethanol (ALCOREX, [42]) (**Fig. 2E**). To identify firing parameters that are modulated by BK channels in mHb neurons, the effect of paxilline (300 nM) was first tested in WT slices. Paxilline significantly reduced spike threshold (q=0.0019), tended to increase spike half-width (q=0.065), and did not affect spike afterhyperpolarization, amplitude, rise time, or fall time (q’s>0.36, **Fig. 2F-I and Suppl. Fig. 1A- C**). The effect of ethanol (50 mM) was then tested in both WT and KI slices. Ethanol reduced spike threshold in WT (q=0.0091) but not in KI (q=0.45) cells (**Fig. 2G**). Ethanol did not significantly affect any of the other parameters in either WT or KI cells (q’s>0.34, **Fig. 2H-I and Suppl. Fig. 1A-C**). In conclusion, ethanol mimics BK channel blockade in mHb neurons and this effect is abolished by the K361N substitution.

### The BK α K361N substitution does not affect sensitivity to acute effects of alcohol in vivo

We first verified that the mutation did not alter the clearance rate of ethanol (effect of time: F_1.202,9.616_=359.6, p<0.0001; effect of genotype: F_2,8_=0.01, p=0.99; time x genotype interaction: F_4,16_=0.2, p=0.91, **Fig. 3A**). In the rotarod assay, WT, Het, and KI males were similarly sensitive to the loss of motor coordination induced by 1.5 g/kg ethanol; there was no effect of genotype on ataxia duration (F_2,26_=1.0, p=0.38) and on BECs measured at recovery (F_2,26_=2.0, p=0.16) (**Fig. 3B**). Likewise, WT, Het and KI males exhibited similar durations of loss-of-righting- response following administration of 3.5 g/kg ethanol (F_2,26_=0.5, p=0.95) and similar BECs at recovery (F_2,26_=0.06, p=0.94) (**Fig. 3C**). The amplitude of hypothermia was also identical across genotypes (effect of time: F_1.666,43.31_=239.6, p<0.0001; effect of genotype: F_2,26_=0.4, p=0.66; time x genotype interaction: F_4,52_=0.5, p=0.71, **Fig. 3D**). Ethanol exerted similar analgesic effects in WT, Het and KI males at 1.5-2.5 g/kg doses (effect of dose: F_2.249,44.98_=61.0, p<0.0001; effect of genotype: F_2,20_=2.0, p=0.16; dose x genotype interaction: F_6,60_=0.6, p=0.73, **Fig. 3E**). Finally, the rewarding effect of 2 g/kg ethanol was equivalent in WT and KI males, as measured by conditioned place preference (effect of conditioning: F_1,24_=25.6, p<0.0001; effect of genotype: F_1,24_=0.6, p=0.43; conditioning x genotype interaction: F_1,24_=0.04, p=0.84, **Fig. 3F**). There was no effect of genotype or conditioning on the distance traveled in the apparatus (**Suppl. Fig. 2**). Altogether, we did not detect any influence of the BK α K361N substitution on the sensitivity of male mice to multiple behavioral and physiological acute effects of moderate and high doses of ethanol.

**Figure 3.**
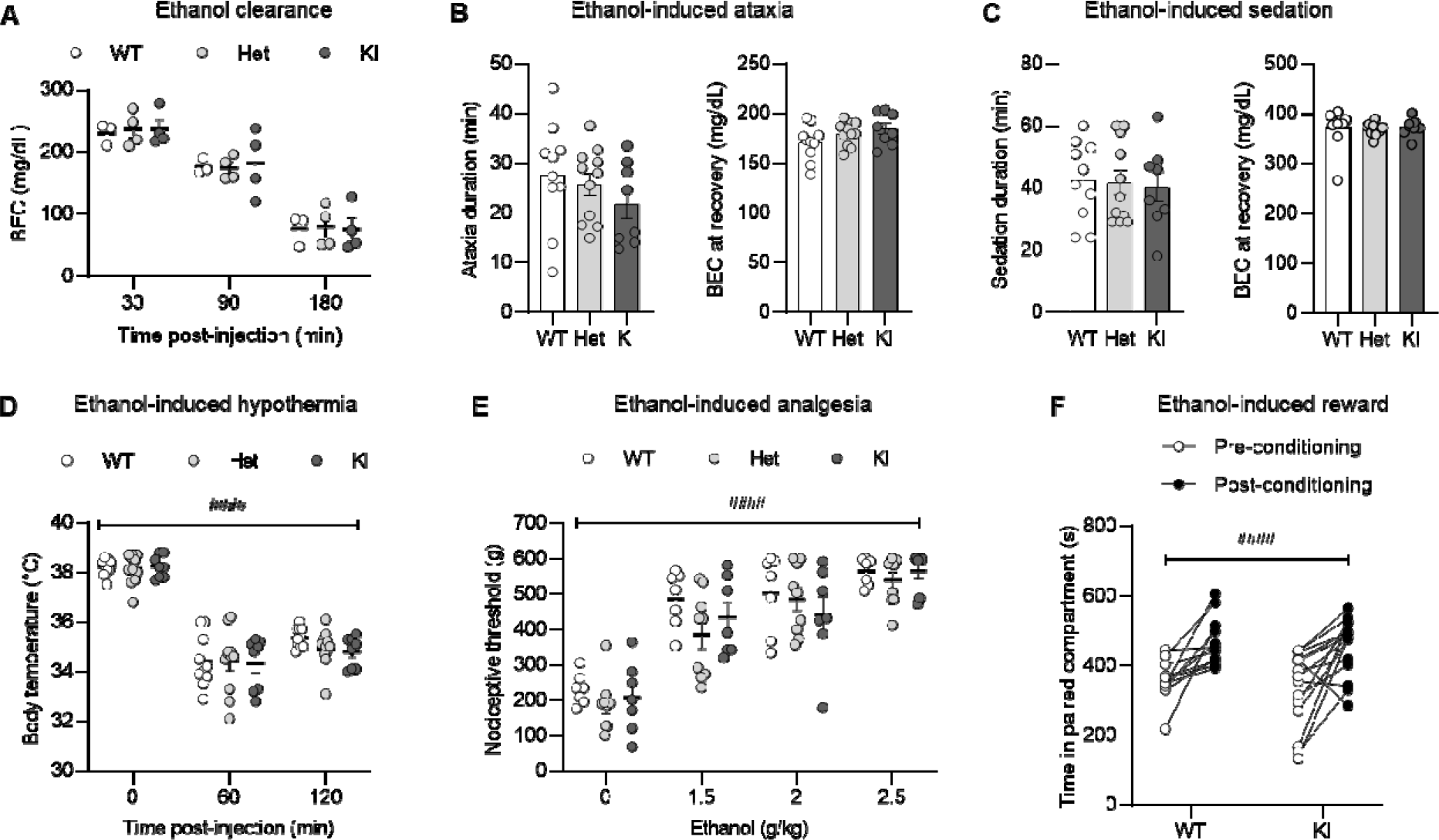
Effect of the BK. α **K361N substitution on acute alcohol sensitivity *in vivo*.** Measures of alcohol metabolism and intoxication were obtained in BK α K361N WT, Het and KI males acutely exposed to ethanol (i.p.). **A.** Blood ethanol concentration (BEC) clearance time- course. **B-F.** Ethanol-induced ataxia (**B**, fixed-speed rotarod), sedation (**C**, loss of righting response), hypothermia (**D**), analgesia (**E**, tail pressure test) and reward (**F**, conditioned place preference). Main effect of ethanol: ^###^, p<0.001. None of the measures was significantly affected by genotype.

### The BK ***α*** K361N substitution does not reproducibly alter alcohol drinking under limited, continuous, or intermittent access

A first group of mice (all males) was given access to voluntary alcohol consumption in 2-h 2BC sessions. The KI males consumed more alcohol than their WT counterparts during the first baseline (BL) week, but the difference subsided by the second week, with the two genotypes stabilizing at similar levels (main effect of genotype, F_1,32_=3.5, p=0.069; *posthoc* on BL1, p=0.032; BL2, p=0.75; **Fig. 4A** and **Suppl. Fig. 3A**). Half of the mice were then exposed to weeks of CIE to trigger voluntary intake escalation during intercalated weeks of 2BC drinking [28] (**Fig. 4B** and **Suppl. Fig. 3B**). As expected, there was a significant week x vapor interaction (F_4,120_=5.1, p=0.0008), reflecting the escalation of voluntary alcohol consumption in CIE mice but not Air mice. A significant main effect of vapor was detected on each post-vapor (PV) week (F_1,30_>7.0, p<0.02), along with a trend for genotype effect on PV4 (F_1,30_=2.9, p=0.097), whereby KI mice consumed less alcohol than WT mice. To follow up on the BL1 and PV4 observations, we repeated the experiment and included both sexes (**Fig. 4C-F** and **Suppl. Fig. 3C-F**). In this second experiment, there was no effect of genotype or trend thereof on any of the baseline or post-vapor weeks. A significant main effect of vapor was detected at PV2, PV3, and PV4 in males (F_1,37_>6.9, p<0.02), and PV3 and PV4 in females (F_1,39_>7.5, p<0.01).

**Figure 4.**
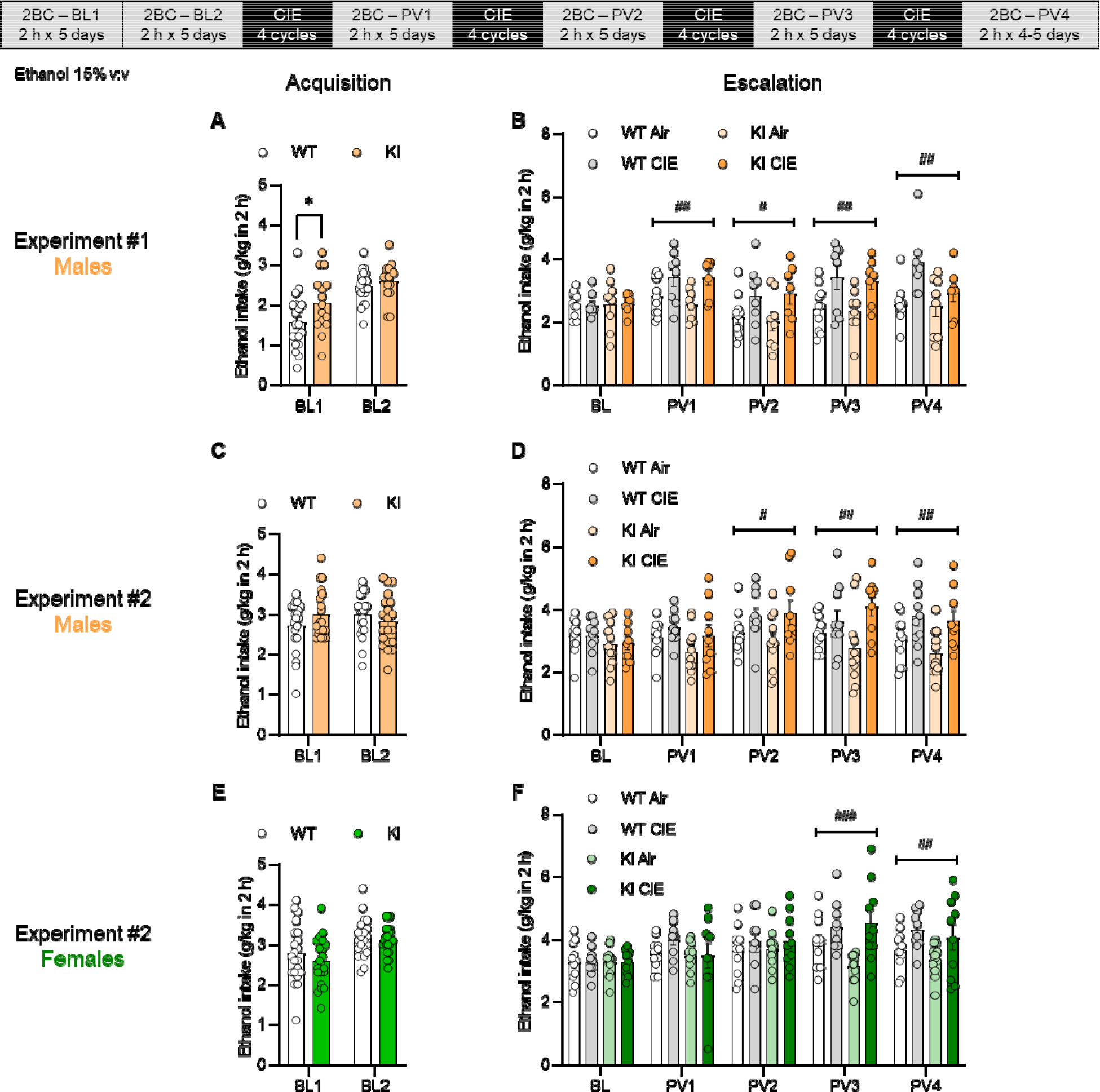
Effect of the BK. α **K361N substitution on baseline and escalated alcohol drinking in the CIE-2BC model.** BK α K361N WT and KI mice were given access to voluntary alcohol consumption in 2-h two-bottle choice sessions prior to (Acquisition, **A**, **C**, **E**) and in-between (Escalation, **B**, **D**, **F**) weeks of chronic intermittent ethanol (CIE) vapor inhalation. A first experiment was conducted in males only (**A-B**) and a repeat experiment included both males (**C-D**) and females (**E-F**). Weekly averages are shown (daily values are provided in **Suppl.** Fig. 3). *, p<0.05, WT *vs.* KI. Main effect of vapor: ^#^, p<0.05; ^##^, p<0.01; ^###^, p<0.001.

BK α WT and K361N KI mice were then tested in another model of alcohol drinking escalation in which access to alcohol switches from continuous to intermittent for 24-h periods [29, 30]. As expected, intermittent access significantly increased both ethanol intake and ethanol preference in both males (intake: F_1,10_=22.9, p=0.0007; preference: F_1,10_=12.8, p=0.0050, **Fig. 5A** and **Suppl. Fig. 4A-B**) and females (intake: F_1,10_=14.9, p=0.0032; preference: F_1,10_=32.3, p=0.0002, **Fig. 5B** and **Suppl. Fig. 4C-D**). In addition, a significant main effect of genotype was detected in females, whereby KI mice had a lower preference for ethanol than their WT counterparts (F_1,10_=5.4, p=0.042, **Fig. 5B**). Finally, there was no effect of genotype on ethanol intake and BECs measured 1 h after alcohol access resumption (**Fig. 5C**). As expected, BECs correlated significantly with intake in both genotypes (WT: R^2^=0.68, p=0.0019; KI: R^2^=0.83, p<0.0001). To follow up on the preference phenotype, we repeated the experiment in a separate group of females. Intermittent access again increased ethanol intake (F_1,15_=28.8, p<0.0001) and preference (F_1,15_=19.4, p=0.0005), but there was no effect of genotype or trend thereof on either measure (**Suppl. Fig. 5**).

**Figure 5.**
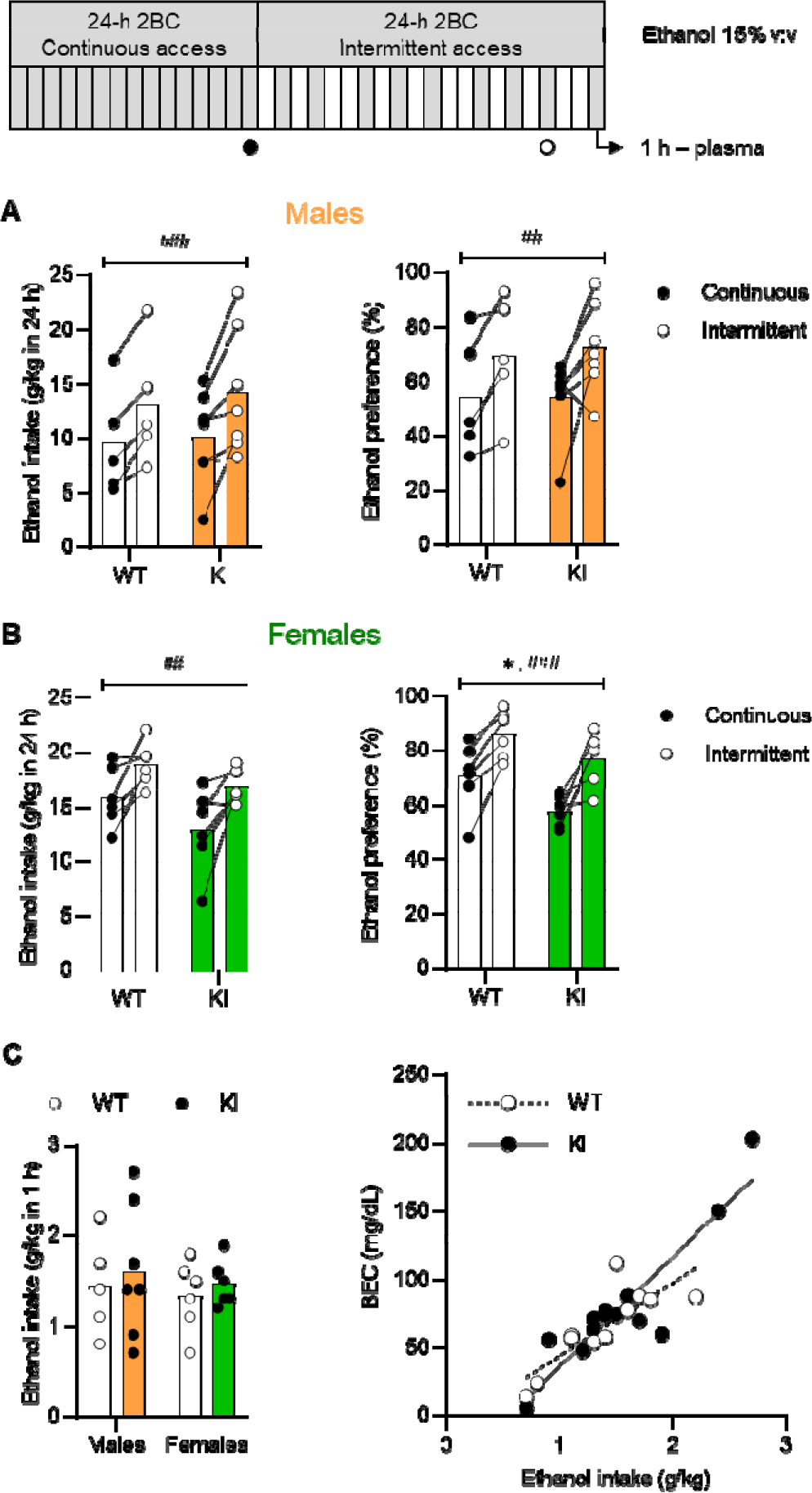
Effect of the BK. α **K361N substitution on alcohol drinking under continuous and intermittent access.** BK α K361N WT and KI males (**A**) and females (**B**) were given continuous access to voluntary alcohol consumption (two-bottle choice) before switching to an intermittent schedule of access (24 h, three times per week). Ethanol intake (left) and preference (right) are shown for the last 24-h period of each phase (daily values are provided in **Suppl.** Fig. 4). Main effect of access schedule: ^##^, p<0.01; ^###^, p<0.001. Main effect of genotype: *, p<0.05. **C.** On the last day of intermittent access, blood ethanol concentrations were measured 1 h after alcohol introduction to verify correlation with intake.

In conclusion, the BK α K361N substitution does not reliably affect alcohol drinking under multiple modalities of moderate and excessive intake.

### The BK ***α*** K361N substitution does not impact the effects of CIE on metabolism, food intake, and locomotor activity

We then tested whether KI mice might exhibit differential sensitivity to other consequences of CIE exposure relevant to AUD, such as metabolic [43] and sleep [44] disturbances. EchoMRI analysis indicated that CIE significantly altered body composition, reducing fat content (F_1,10_=9.8, p=0.011) while increasing lean (F_1,10_=10.6, p=0.0086) and water (F_1,10_=13.8, p=0.0040) content, in the absence of body weight change (F_1,10_=0.001, p=0.98, **Fig. 6A-B**).

**Figure 6.**
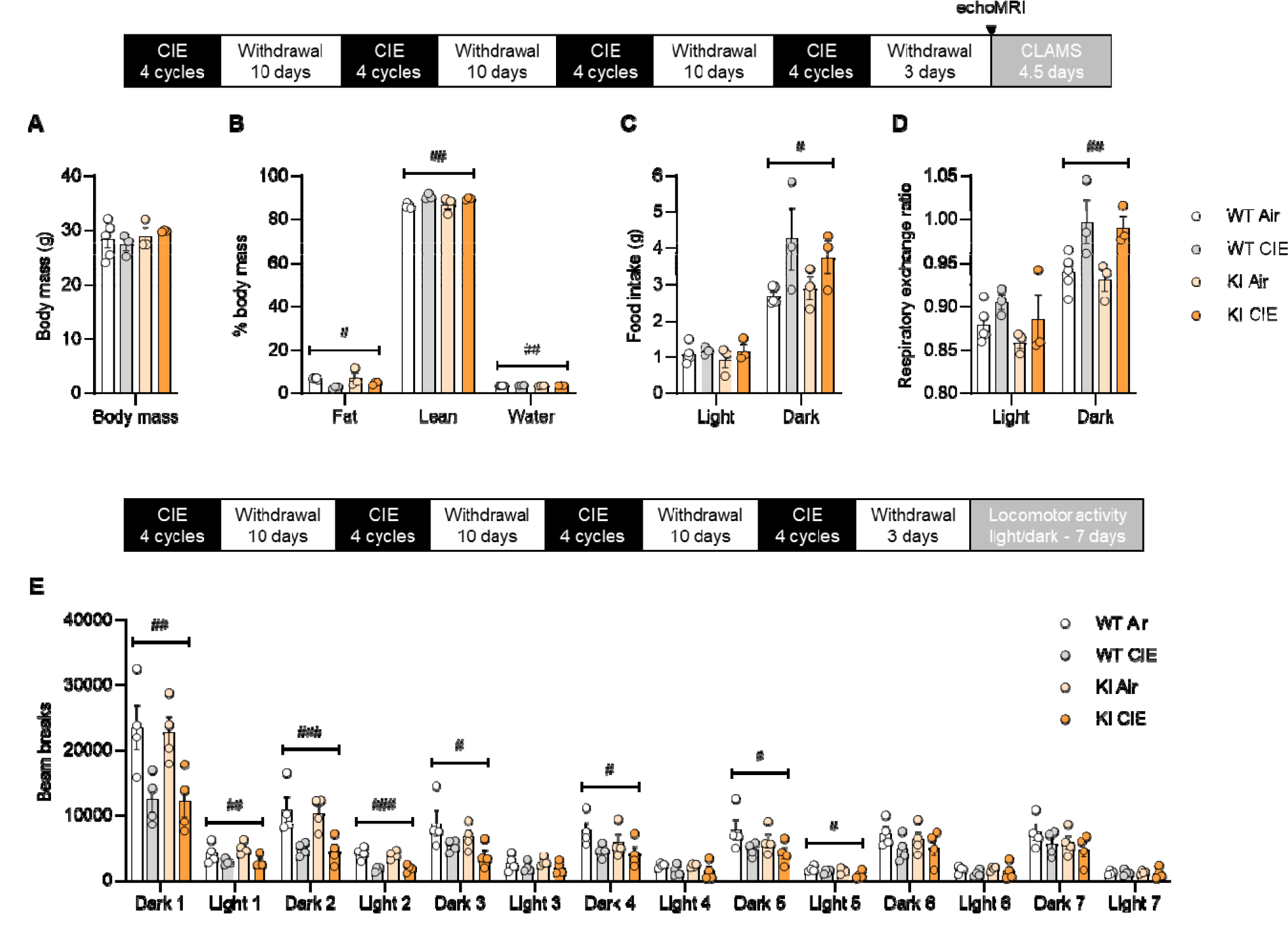
Effect of the BK. α **K361N substitution on the metabolic and locomotor effects of chronic intermittent ethanol (CIE) exposure.** BK α K361N WT and KI males were exposed to air or chronic intermittent ethanol (CIE) vapor inhalation. In a first experiment, body mass (**A**) and body composition (**B**) were determined 3 days into withdrawal. Food intake (**C**) and respiratory exchange ratio (**D**) were then recorded in metabolic chambers during 4.5 days. In a second experiment, locomotor activity was recorded starting 3 days into withdrawal (**E**). Main effect of vapor: ^#^, p<0.05; ^##^, p<0.01; ^###^, p<0.001. None of the measures was significantly affected by genotype.

Metabolic monitoring also revealed increases in dark-phase food intake (F_1,10_=7.3, p=0.023, **Fig. 6C**) and dark-phase RER (F_1,10_=15.7, p=0.0027, **Fig. 6D**) in CIE-withdrawn mice. The K361N substitution did not influence any of these outcomes (F’s<1.0, p’s>0.34 for main effect of genotype and genotype x vapor interaction). Furthermore, neither genotype nor CIE affected actimetry-based sleep measures (**Suppl. Fig. 6**).

Finally, given the role of BK channels in regulating neuronal excitability in the suprachiasmatic nucleus (the primary circadian pacemaker in mammals) [45, 46] and the desynchronization of biological rhythms observed in AUD [47, 48], we sought to determine whether the action of ethanol on BK channels could be responsible for a disruption of circadian rhythmicity in CIE-exposed mice. Under a standard light-dark cycle, the ambulation of CIE-exposed mice was significantly reduced up to withdrawal day 8 (vapor x time interaction: F_13,156_=10.3, p<0.0001, see **Fig. 6E** for significance of vapor effect at individual timepoints). There was no significant influence of genotype on ambulation nor on the depressant effect of CIE withdrawal (genotype effect: F_1,12_=0.5, p=0.49; genotype x vapor interaction: F_1,12_=0.08, p=0.78). To test the function of the intrinsic pacemaker, mice were then switched to constant darkness and chi-square periodogram analysis was used to determine the period length and relative power of the dominant circadian component of ambulation counts (**Suppl. Fig. 7**). Two-way ANOVA revealed a significant interaction between vapor and genotype on period length (F_1,12_=6.5, p=0.025), but none of the pairwise comparisons reached statistical significance. Neither the K361N substitution nor alcohol withdrawal significantly affected the relative power (genotype effect: F_1,12_=2.5, p=0.14; vapor effect: F_1,12_=2.2, p=0.17; genotype x vapor interaction: F_1,12_=0.60, p=0.45).

## Discussion

Altogether, our data demonstrate that BK channel inhibition can modulate alcohol drinking in mice, but the underlying mechanism does not involve the interaction of ethanol with BK α residue K361. This interaction is also not required for alcohol to produce ataxia, sedation, hypothermia, analgesia, and reward, nor for CIE to produce metabolic and locomotor changes during withdrawal.

The major behavioral disturbances elicited by the blockade of BK channels have historically been a hurdle to analyze the behavioral relevance of ethanol’s action at this target. This limitation is illustrated by the results of our pharmacological experiments, whereby the dose-dependent effects of penitrem A on ethanol intake had to be disentangled from tremor induction. The dose of 0.1 mg/kg reduced alcohol drinking both at baseline and following additional weeks of 2BC in Air mice, without inducing significant tremor or affecting saccharin drinking. The insensitivity of CIE mice to this dose may reflect adaptive processes producing molecular tolerance to ethanol [12, 42, 49–51] or a possible upregulation of BK channels by CIE, as seen in alcohol-exposed flies [9–11]. In contrast, the dose of 0.05 mg/kg increased alcohol intake.

This effect was only detected after additional weeks of 2BC in Air mice, suggesting that it requires long-term alcohol drinking and may have been masked by intake escalation in CIE mice (ceiling effect). The dose-dependent pattern of penitrem A’s effects on alcohol drinking may reflect gradually insurmountable blockade (with ethanol acting as a BK channel activator) or priming at 0.05 mg/kg and substitution for ethanol at 0.1 mg/kg (with ethanol acting as a BK channel inhibitor).

Paxilline injected at doses at least ten times lower than doses typically used to induce tremors (6-8 mg/kg, [40, 52]) did not cause overt behavioral abnormalities and did not alter ethanol intake. An even lower dose of paxilline had been previously shown to reverse picrotoxin- and pentylenetetrazole-induced seizures in the absence of tremors [16], which suggests that the doses we used were high enough to significantly reach and block BK channels in the mouse brain, but we cannot rule out that higher doses would have replicated the effects of penitrem A. It is also possible that penitrem A exerted its behavioral effects via a different target that was spared by paxilline, such as GABA_A_ receptors [53]. Consistent with this scenario, BK channel activation by BMS-204352, at a dose known to acutely reverse the sensory hypersensitivity and social interaction deficits of *Fmr1* KO mice [17, 18], had no effect on ethanol intake.

The mHb is a known hotspot for the expression of BK channel α, β4, and γ3 subunits in the brain [54–56]. Using oligoprobes specific for BK α alternatively spliced exons STREX and ALCOREX, the mHb was the mouse brain area showing the strongest signal. The presence of β4 and ALCOREX both confer ethanol sensitivity to BK channels [42, 57], making the mHb well-suited for the validation of the K361N substitution. To the best of our knowledge, the effects of paxilline or ethanol on mHb BK channels had never been reported. We found that inhibition of mHb BK channels by paxilline hyperpolarized spike threshold, as previously described in hippocampal granule cells [58]. Spike width and afterhyperpolarization, two parameters regulated by BK channels in other neuronal populations [see 59 for review], had inconsistent sensitivity to paxilline in WT mHb neurons. Ethanol’s effects in WT slices were similar to those of paxilline (reduced spike threshold, no effect on other parameters), suggesting that ethanol may inhibit BK channels in the mHb, as previously suggested in the central amygdala [36]. The lack of effect of ethanol on spike threshold in KI slices supports a direct mechanism and confirms the resistance of K361N channels to ethanol in a native setting.

Preventing ethanol from interacting with BK α via the K361N substitution did not significantly alter the acute behavioral responses to alcohol examined here in mice. This finding contrasts with the ability of a genetic manipulation in worms (T381I substitution) to reduce the effects of acute ethanol on egg laying and locomotion [7]. The homologous mammalian residue (T352) is adjacent to the alcohol-sensing site identified by Bukiya *et al.* [15], but the effect of mutating this residue remains to be tested in mice.

The lack of a consistent effect of K361N on CIE-induced ethanol intake escalation is also at odds with the phenotypes of BK β1 and BK β4 knockout mice we had reported in this paradigm [13]. Based on the known influence of BK β1 and β4 subunits on the potentiation of BK-mediated currents by ethanol [57], our previous results suggested that the action of ethanol at BK channels promotes escalation. We thus expected that the K361N substitution would hinder escalation. The negative outcome reported here suggests that the influence of BK channels on alcohol drinking might be indirect (e.g., recruitment of BK channels downstream of another primary target of ethanol). This scenario would be compatible with intact alcohol drinking in KI mice, given that the K361N substitution does not impair the physiological activity of BK channels [15]. Alternatively, the role of BK channels in the regulation of alcohol drinking may be mediated by BK channel auxiliary subunits and/or by a BK α site(s) different from K361. This scenario is relevant to alcohol-induced constriction of cerebral arteries, which is mediated by the inhibition of myocyte BK channels and requires BK β1 [60] but is unchanged in BK α K361N KI mice (Kuntamallappanavar G, Bukiya AN, and Dopico AM; unpublished results).

We hypothesized that, aside from alcohol drinking escalation, ethanol’s action at BK channels may mediate other physiological consequences of CIE exposure. We detected significant effects of CIE on metabolism, which, to the best of our knowledge, had never been reported in this mouse model. We found that 4 weeks of CIE significantly altered the body composition of mice, reducing fat content and increasing lean content without affecting their total body mass. This observation is consistent with reports of reduced body fat in chronic alcoholics, in the absence of body weight change and in proportion to the level of alcohol consumption [61–64]. Studies in mice chronically fed an alcohol liquid diet have indicated that chronic alcohol reduces white, rather than brown, adipose tissue and that such lipolysis is associated with hepatic steatosis, i.e. ectopic deposition of fat in the liver [see 43 for review]. Interestingly, CIE-exposed rats and mice do not show evidence of hepatic steatosis [65, 66]. The CIE procedure may therefore induce changes in lipid metabolism that reflect an early stage of the development of alcohol liver disease. Multiple molecular mechanisms have been proposed to underlie alcohol-induced lipolysis [reviewed in 43]; our data indicate that the interaction of ethanol with BK α K361 is not involved in this phenomenon.

The leaner phenotype of CIE-exposed mice was associated with a significant increase in food intake during the first week of withdrawal, which may reflect a homeostatic adaptation to the loss of body fat. In humans, chronic alcohol abuse increases daily caloric intake, yet alcohol represents a substantial fraction of this intake, such that energy intake provided only by food ingestion is typically lower than in healthy counterparts [61, 62, 67]. In one study, 14 days of abstinence normalized the nutritional status of the AUD subjects, but it is not known whether a compensatory increase in food intake may have occurred during their first week of abstinence [67]. Withdrawal from CIE was also associated with a robust increase in RER, reflecting preferential utilization of carbohydrates as a fuel. This pattern may result from deficient lipid storage, as reflected by reduced body fat, and a corresponding inability to sustain normal levels of fatty acid oxidation. However, this observation contrasts with the lower respiratory quotient, higher lipid oxidation, and reduced carbohydrate oxidation reported in human alcoholics, which all normalize after three months of abstinence [61, 62, 67, 68]. To the best of our knowledge, the possibility that a rebound increase in respiratory quotient may occur during the first week of abstinence has not been explored in humans. In any case, the phenotype of KI mice indicates that the action of ethanol at BK α K361 is not responsible for the metabolic and nutritional changes associated with early withdrawal from chronic alcohol exposure.

One limitation of the pharmacological and genetic manipulations used in the present study is that they were systemic and constitutive, respectively, which may have occluded phenotypic changes if BK channels expressed in different brain regions exert opposing effects on a given alcohol-driven behavior. Further complicating the situation, ethanol has differential effects on BK channel activity in different neuronal populations, owing at least in part to the complement of auxiliary subunits expressed by these cells: activation in neurohypophysial terminals [1, 35], nucleus accumbens somas [2], and external globus pallidus Npas1 neurons [3]; putative inhibition in central amygdala terminals [36] and mHb somas (this study); no effect in supraoptic nucleus cell bodies [35] and nucleus accumbens dendrites [2]. These region- and subcellular compartment-specific subpopulations of BK channels may in turn be differentially sensitive to the effects of pharmacological modulators. Accordingly, the effect of a ligand or mutation in a given brain region or neural circuit may offset its effect elsewhere. Another important consideration is that the present investigation focused on effects of alcohol that we hypothesized to be mediated by neuronal BK channels. However, BK channels are also expressed by other brain cell types, with particularly high levels found in astrocytes and oligodendrocyte precursor cells [59, 69, 70]. Future studies will aim to determine whether alcohol’s effects on the function of glial cells might depend on the interaction of ethanol with BK α K361.

In conclusion, our data show that, in the mouse, ethanol’s interaction with BK α K361 does not mediate the role of BK channels in the modulation of moderate and excessive alcohol drinking. This interaction is also not required for acute responses to various doses of ethanol or for the metabolic consequences of CIE exposure. It remains possible, however, that the action of ethanol at BK α K361 mediates molecular, cellular, or behavioral effects of acute or chronic alcohol that were not examined in the present study.

## Acknowledgments

We thank Dr. Elizabeth Thomas for lending us her rotarod apparatus, Dr. Mark Azar for his help with the tremor experiment, Dr. Fu-Ming Zhou for his assistance with electrophysiological validation of the BK α K361N substitution, Brandon Hedges for his assistance with *in vivo* paxilline testing, and Carolyn Ferguson for expert technical assistance in the generation of knockin mice. We are also grateful for the support of TSRI’s Animal Models Core, the Integrative Neuroscience Initiative on Alcoholism-Neuroimmune, and the TSRI Alcohol Research Center Animal Models Core, which conducted BEC analysis for this study.

## Data availability

All data supporting the findings of this study are available within the article and its supplementary information files.

## Author contributions

AO: investigation, methodology, formal analysis. MK: investigation, methodology, software. PJG: Validation, formal analysis. CL: Investigation. BM: Investigation. MB: Validation, formal analysis. PB: software, formal analysis. AMD: Conceptualization. MR: Methodology, supervision. AJR: Investigation, methodology, formal analysis, supervision. GEH: Methodology, resources, supervision. CC: Conceptualization, funding acquisition, project administration, investigation, formal analysis, supervision, writing.

## References

1. Dopico AM, Lemos JR, and Treistman SN, Ethanol increases the activity of large conductance, Ca(2+)-activated K+ channels in isolated neurohypophysial terminals. Mol Pharmacol, 1996. 49(1):40–8.

2. Martin G, Puig S, Pietrzykowski A, Zadek P, Emery P, and Treistman S, Somatic localization of a specific large-conductance calcium-activated potassium channel subtype controls compartmentalized ethanol sensitivity in the nucleus accumbens. J Neurosci, 2004. 24(29):6563–72.

3. Abrahao KP, Chancey JH, Chan CS, and Lovinger DM, Ethanol-Sensitive Pacemaker Neurons in the Mouse External Globus Pallidus. Neuropsychopharmacology, 2017. 42(5):1070–1081.

4. Dopico AM, Bukiya AN, Kuntamallappanavar G, and Liu J, Modulation of BK Channels by Ethanol. Int Rev Neurobiol, 2016. 128:239–79.

5. Bettinger JC and Davies AG, The role of the BK channel in ethanol response behaviors: evidence from model organism and human studies. Front Physiol, 2014. 5:346.

6. Davies AG, Pierce-Shimomura JT, Kim H, VanHoven MK, Thiele TR, Bonci A, Bargmann CI, and McIntire SL, A central role of the BK potassium channel in behavioral responses to ethanol in C. elegans. Cell, 2003. 115(6):655–66.

7. Davis SJ, Scott LL, Hu K, and Pierce-Shimomura JT, Conserved single residue in the BK potassium channel required for activation by alcohol and intoxication in C. elegans. J Neurosci, 2014. 34(29):9562–73.

8. Cowmeadow RB, Krishnan HR, and Atkinson NS, The slowpoke gene is necessary for rapid ethanol tolerance in Drosophila. Alcohol Clin Exp Res, 2005. 29(10):1777–86.

9. Cowmeadow RB, Krishnan HR, Ghezzi A, Al’Hasan YM, Wang YZ, and Atkinson NS, Ethanol tolerance caused by slowpoke induction in Drosophila. Alcohol Clin Exp Res, 2006. 30(5):745–53.

10. Ghezzi A, Pohl JB, Wang Y, and Atkinson NS, BK channels play a counter-adaptive role in drug tolerance and dependence. Proc Natl Acad Sci U S A, 2010. 107(37):16360–5.

11. Ghezzi A, Al-Hasan YM, Larios LE, Bohm RA, and Atkinson NS, slo K(+) channel gene regulation mediates rapid drug tolerance. Proc Natl Acad Sci U S A, 2004. 101(49):17276–81.

12. Martin GE, Hendrickson LM, Penta KL, Friesen RM, Pietrzykowski AZ, Tapper AR, and Treistman SN, Identification of a BK channel auxiliary protein controlling molecular and behavioral tolerance to alcohol. Proc Natl Acad Sci U S A, 2008. 105(45):17543–8.

13. Kreifeldt M, Le D, Treistman SN, Koob GF, and Contet C, BK channel beta1 and beta4 auxiliary subunits exert opposite influences on escalated ethanol drinking in dependent mice. Front Integr Neurosci, 2013. 7:105.

14. Kreifeldt M, Cates-Gatto C, Roberts AJ, and Contet C, BK Channel beta1 Subunit Contributes to Behavioral Adaptations Elicited by Chronic Intermittent Ethanol Exposure. Alcohol Clin Exp Res, 2015. 39(12):2394–402.

15. Bukiya AN, Kuntamallappanavar G, Edwards J, Singh AK, Shivakumar B, and Dopico AM, An alcohol-sensing site in the calcium- and voltage-gated, large conductance potassium (BK) channel. Proc Natl Acad Sci U S A, 2014. 111(25):9313–8.

16. Sheehan JJ, Benedetti BL, and Barth AL, Anticonvulsant effects of the BK-channel antagonist paxilline. Epilepsia, 2009. 50(4):711–20.

17. Zhang Y, Bonnan A, Bony G, Ferezou I, Pietropaolo S, Ginger M, Sans N, Rossier J, Oostra B, LeMasson G, and Frick A, Dendritic channelopathies contribute to neocortical and sensory hyperexcitability in Fmr1(-/y) mice. Nat Neurosci, 2014. 17(12):1701–9.

18. Hebert B, Pietropaolo S, Meme S, Laudier B, Laugeray A, Doisne N, Quartier A, Lefeuvre S, Got L, Cahard D, Laumonnier F, Crusio WE, Pichon J, Menuet A, Perche O, and Briault S, Rescue of fragile X syndrome phenotypes in Fmr1 KO mice by a BKCa channel opener molecule. Orphanet J Rare Dis, 2014. 9:124.

19. Blednov YA, Borghese CM, Ruiz CI, Cullins MA, Da Costa A, Osterndorff-Kahanek EA, Homanics GE, and Harris RA, Mutation of the inhibitory ethanol site in GABAA rho1 receptors promotes tolerance to ethanol-induced motor incoordination. Neuropharmacology, 2017. 123:201–209.

20. Hsu PD, Scott DA, Weinstein JA, Ran FA, Konermann S, Agarwala V, Li Y, Fine EJ, Wu X, Shalem O, Cradick TJ, Marraffini LA, Bao G, and Zhang F, DNA targeting specificity of RNA-guided Cas9 nucleases. Nat Biotechnol, 2013. 31(9):827–32.

21. Bassett AR, Tibbit C, Ponting CP, and Liu JL, Highly Efficient Targeted Mutagenesis of Drosophila with the CRISPR/Cas9 System. Cell Rep, 2014. 6(6):1178–1179.

22. Yang H, Wang H, and Jaenisch R, Generating genetically modified mice using CRISPR/Cas-mediated genome engineering. Nat Protoc, 2014. 9(8):1956–68.

23. Guo D, Li X, Zhu P, Feng Y, Yang J, Zheng Z, Yang W, Zhang E, Zhou S, and Wang H, Online High-throughput Mutagenesis Designer Using Scoring Matrix of Sequence-specific Endonucleases. J Integr Bioinform, 2015. 12(1):35–48.

24. Verheij MMM, Contet C, Karel P, Latour J, van der Doelen RHA, Geenen B, van Hulten JA, Meyer F, Kozicz T, George O, Koob GF, and Homberg JR, Median and Dorsal Raphe Serotonergic Neurons Control Moderate Versus Compulsive Cocaine Intake. Biol Psychiatry, 2018. 83(12):1024–1035.

25. V Szücs A. NeuroExpress program for analyzing patch-clamp data. 2022; Available from: https://www.researchgate.net/publication/338623194_NeuroExpress_program_for_analyzing_patch-clamp_data.

26. Gallagher RT and Hawkes AD, The potent tremorgenic neurotoxins lolitrem B and aflatrem: a comparison of the tremor response in mice. Experientia, 1986. 42(7):823–5.

27. Elhabazi K, Ayachi S, Ilien B, and Simonin F, Assessment of morphine-induced hyperalgesia and analgesic tolerance in mice using thermal and mechanical nociceptive modalities. J Vis Exp, 2014(89):e51264.

28. Becker HC and Lopez MF, Increased ethanol drinking after repeated chronic ethanol exposure and withdrawal experience in C57BL/6 mice. Alcohol Clin Exp Res, 2004. 28(12):1829–38.

29. Melendez RI, Intermittent (every-other-day) drinking induces rapid escalation of ethanol intake and preference in adolescent and adult C57BL/6J mice. Alcohol Clin Exp Res, 2011. 35(4):652–8.

30. Hwa LS, Chu A, Levinson SA, Kayyali TM, DeBold JF, and Miczek KA, Persistent escalation of alcohol drinking in C57BL/6J mice with intermittent access to 20% ethanol. Alcohol Clin Exp Res, 2011. 35(11):1938–47.

31. McLean JA and Tobin G, Animal and Human Calorimetry. 1987, Cambridge: Cambridge University Press. 338.

32. Pack AI, Galante RJ, Maislin G, Cater J, Metaxas D, Lu S, Zhang L, Von Smith R, Kay T, Lian J, Svenson K, and Peters LL, Novel method for high-throughput phenotyping of sleep in mice. Physiol Genomics, 2007. 28(2):232–8.

33. Enright JT, The search for rhythmicity in biological time-series. J Theor Biol, 1965. 8(3):426–68.

34. Refinetti R, Laboratory instrumentation and computing: comparison of six methods for the determination of the period of circadian rhythms. Physiol Behav, 1993. 54(5):869–75.

35. Dopico AM, Widmer H, Wang G, Lemos JR, and Treistman SN, Rat supraoptic magnocellular neurones show distinct large conductance, Ca2+-activated K+ channel subtypes in cell bodies versus nerve endings. J Physiol, 1999. 519 Pt 1(Pt 1):101–14.

36. Li Q, Madison R, and Moore SD, Presynaptic BK channels modulate ethanol-induced enhancement of GABAergic transmission in the rat central amygdala nucleus. J Neurosci, 2014. 34(41):13714–24.

37. Knaus HG, McManus OB, Lee SH, Schmalhofer WA, Garcia-Calvo M, Helms LM, Sanchez M, Giangiacomo K, Reuben JP, Smith AB, 3rd, and et al., Tremorgenic indole alkaloids potently inhibit smooth muscle high-conductance calcium-activated potassium channels. Biochemistry, 1994. 33(19):5819–28.

38. Asano S, Bratz IN, Berwick ZC, Fancher IS, Tune JD, and Dick GM, Penitrem A as a tool for understanding the role of large conductance Ca(2+)/voltage-sensitive K(+) channels in vascular function. J Pharmacol Exp Ther, 2012. 342(2):453–60.

39. Jortner BS, Ehrich M, Katherman AE, Huckle WR, and Carter ME, Effects of prolonged tremor due to penitrem A in mice. Drug Chem Toxicol, 1986. 9(2):101–16.

40. Imlach WL, Finch SC, Dunlop J, Meredith AL, Aldrich RW, and Dalziel JE, The molecular mechanism of “ryegrass staggers,” a neurological disorder of K+ channels. J Pharmacol Exp Ther, 2008. 327(3):657–64.

41. Sausbier M, Hu H, Arntz C, Feil S, Kamm S, Adelsberger H, Sausbier U, Sailer CA, Feil R, Hofmann F, Korth M, Shipston MJ, Knaus HG, Wolfer DP, Pedroarena CM, Storm JF, and Ruth P, Cerebellar ataxia and Purkinje cell dysfunction caused by Ca2+-activated K+ channel deficiency. Proc Natl Acad Sci U S A, 2004. 101(25):9474–8.

42. Pietrzykowski AZ, Friesen RM, Martin GE, Puig SI, Nowak CL, Wynne PM, Siegelmann HT, and Treistman SN, Posttranscriptional regulation of BK channel splice variant stability by miR-9 underlies neuroadaptation to alcohol. Neuron, 2008. 59(2):274–87.

43. Steiner JL and Lang CH, Alcohol, Adipose Tissue and Lipid Dysregulation. Biomolecules, 2017. 7(1).

44. Chakravorty S, Chaudhary NS, and Brower KJ, Alcohol Dependence and Its Relationship With Insomnia and Other Sleep Disorders. Alcohol Clin Exp Res, 2016. 40(11):2271–2282.

45. Meredith AL, Wiler SW, Miller BH, Takahashi JS, Fodor AA, Ruby NF, and Aldrich RW, BK calcium-activated potassium channels regulate circadian behavioral rhythms and pacemaker output. Nat Neurosci, 2006. 9(8):1041–9.

46. Harvey JRM, Plante AE, and Meredith AL, Ion Channels Controlling Circadian Rhythms in Suprachiasmatic Nucleus Excitability. Physiol Rev, 2020. 100(4):1415–1454.

47. Lindberg D, Andres-Beck L, Jia YF, Kang S, and Choi DS, Purinergic Signaling in Neuron-Astrocyte Interactions, Circadian Rhythms, and Alcohol Use Disorder. Front Physiol, 2018. 9:9.

48. Meyrel M, Rolland B, and Geoffroy PA, Alterations in circadian rhythms following alcohol use: A systematic review. Prog Neuropsychopharmacol Biol Psychiatry, 2020. 99:109831.

49. Pietrzykowski AZ, Martin GE, Puig SI, Knott TK, Lemos JR, and Treistman SN, Alcohol tolerance in large-conductance, calcium-activated potassium channels of CNS terminals is intrinsic and includes two components: decreased ethanol potentiation and decreased channel density. J Neurosci, 2004. 24(38):8322–32.

50. Velazquez-Marrero C, Wynne P, Bernardo A, Palacio S, Martin G, and Treistman SN, The relationship between duration of initial alcohol exposure and persistence of molecular tolerance is markedly nonlinear. J Neurosci, 2011. 31(7):2436–46.

51. Palacio S, Velazquez-Marrero C, Marrero HG, Seale GE, Yudowski GA, and Treistman SN, Time-Dependent Effects of Ethanol on BK Channel Expression and Trafficking in Hippocampal Neurons. Alcohol Clin Exp Res, 2015. 39(9):1619–31.

52. Combs MD, Hamlin A, and Quinn JC, A single exposure to the tremorgenic mycotoxin lolitrem B inhibits voluntary motor activity and spatial orientation but not spatial learning or memory in mice. Toxicon, 2019. 168:58–66.

53. Moldes-Anaya AS, Fonnum F, Eriksen GS, Rundberget T, Walaas SI, and Wigestrand MB, In vitro neuropharmacological evaluation of penitrem-induced tremorgenic syndromes: importance of the GABAergic system. Neurochem Int, 2011. 59(7):1074–81.

54. Chang CP, Dworetzky SI, Wang J, and Goldstein ME, Differential expression of the alpha and beta subunits of the large-conductance calcium-activated potassium channel: implication for channel diversity. Brain Res Mol Brain Res, 1997. 45(1):33–40.

55. Zhang YY, Han X, Liu Y, Chen J, Hua L, Ma Q, Huang YY, Tang QY, and Zhang Z, +mRNA expression of LRRC55 protein (leucine-rich repeat-containing protein 55) in the adult mouse brain. PLoS One, 2018. 13(1):e0191749.

56. Allen Institute for Brain Science; Allen Mouse Brain Atlas.

57. Kuntamallappanavar G and Dopico AM, Alcohol modulation of BK channel gating depends on beta subunit composition. J Gen Physiol, 2016. 148(5):419–440.

58. Whitmire LE, Ling L, Bugay V, Carver CM, Timilsina S, Chuang HH, Jaffe DB, Shapiro MS, Cavazos JE, and Brenner R, Downregulation of KCNMB4 expression and changes in BK channel subtype in hippocampal granule neurons following seizure activity. PLoS One, 2017. 12(11):e0188064.

59. Contet C, Goulding SP, Kuljis DA, and Barth AL, BK Channels in the Central Nervous System. Int Rev Neurobiol, 2016. 128:281–342.

60. Bukiya AN, Liu J, and Dopico AM, The BK channel accessory beta1 subunit determines alcohol-induced cerebrovascular constriction. FEBS Lett, 2009. 583(17):2779–84.

61. Addolorato G, Capristo E, Greco AV, Stefanini GF, and Gasbarrini G, Energy expenditure, substrate oxidation, and body composition in subjects with chronic alcoholism: new findings from metabolic assessment. Alcohol Clin Exp Res, 1997. 21(6):962–7.

62. Addolorato G, Capristo E, Greco AV, Stefanini GF, and Gasbarrini G, Influence of chronic alcohol abuse on body weight and energy metabolism: is excess ethanol consumption a risk factor for obesity or malnutrition? J Intern Med, 1998. 244(5):387–95.

63. Addolorato G, Capristo E, Marini M, Santini P, Scognamiglio U, Attilia ML, Messineo D, Sasso GF, Gasbarrini G, and Ceccanti M, Body composition changes induced by chronic ethanol abuse: evaluation by dual energy X-ray absorptiometry. Am J Gastroenterol, 2000. 95(9):2323–7.

64. Liangpunsakul S, Crabb DW, and Qi R, Relationship among alcohol intake, body fat, and physical activity: a population-based study. Ann Epidemiol, 2010. 20(9):670–5.

65. Mouton AJ, Maxi JK, Souza-Smith F, Bagby GJ, Gilpin NW, Molina PE, and Gardner JD, Alcohol Vapor Inhalation as a Model of Alcohol-Induced Organ Disease. Alcohol Clin Exp Res, 2016. 40(8):1671–8.

66. Xu J, Ma HY, Liu X, Rosenthal S, Baglieri J, McCubbin R, Sun M, Koyama Y, Geoffroy CG, Saijo K, Shang L, Nishio T, Maricic I, Kreifeldt M, Kusumanchi P, Roberts A, Zheng B, Kumar V, Zengler K, Pizzo DP, Hosseini M, Contet C, Glass CK, Liangpunsakul S, Tsukamoto H, Gao B, Karin M, Brenner DA, Koob GF, and Kisseleva T, Blockade of IL- 17 signaling reverses alcohol-induced liver injury and excessive alcohol drinking in mice. JCI Insight, 2020. 5(3).

67. Levine JA, Harris MM, and Morgan MY, Energy expenditure in chronic alcohol abuse. Eur J Clin Invest, 2000. 30(9):779–86.

68. Addolorato G, Capristo E, Greco AV, Caputo F, Stefanini GF, and Gasbarrini G, Three months of abstinence from alcohol normalizes energy expenditure and substrate oxidation in alcoholics: a longitudinal study. Am J Gastroenterol, 1998. 93(12):2476–81.

69. Zhang Y, Chen K, Sloan SA, Bennett ML, Scholze AR, O’Keeffe S, Phatnani HP, Guarnieri P, Caneda C, Ruderisch N, Deng S, Liddelow SA, Zhang C, Daneman R, Maniatis T, Barres BA, and Wu JQ, An RNA-sequencing transcriptome and splicing database of glia, neurons, and vascular cells of the cerebral cortex. J Neurosci, 2014. 34(36):11929–47.

70. Buttigieg J, Eftekharpour E, Karimi-Abdolrezaee S, and Fehlings MG, Molecular and electrophysiological evidence for the expression of BK channels in oligodendroglial precursor cells. Eur J Neurosci, 2011. 34(4):538–47.

