## Supplementary material for "Ethanol’s interaction with BK channel α subunit residue K361 does not mediate behavioral responses to alcohol in mice": Suppl

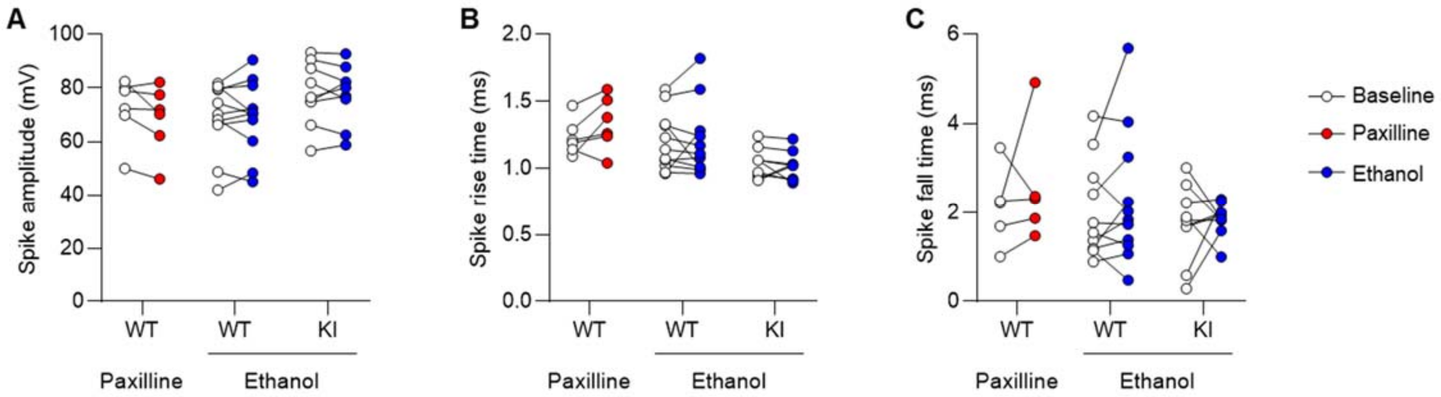


**Supplementary Figure 1. Spike properties of wild-type (WT) and BK α K361N knockin (KI) medial habenula neurons.** Recordings were obtained before (open circles) and during (closed circles) application of the BK channel blocker paxilline (300 nM, red) or ethanol (50 mM, blue). **A.** Spike amplitude. **B.** Spike rise time. **C.** Spike fall time. Paxilline and ethanol had no significant effect on any of the measures.

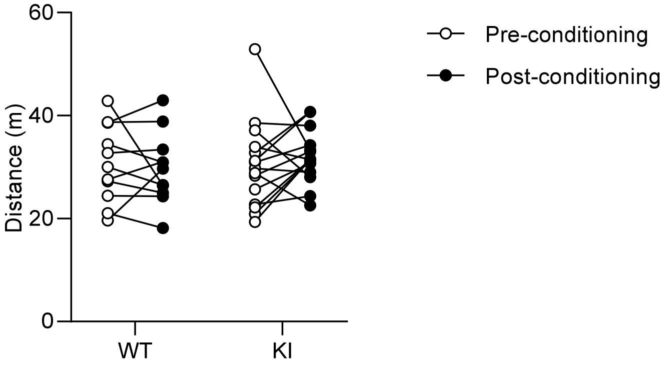


**Supplementary Figure 2. Distance traveled by wild-type (WT) and BK α K361N knockin (KI) mice in the conditioned place preference apparatus.** There was no significant effect of genotype or conditioning.

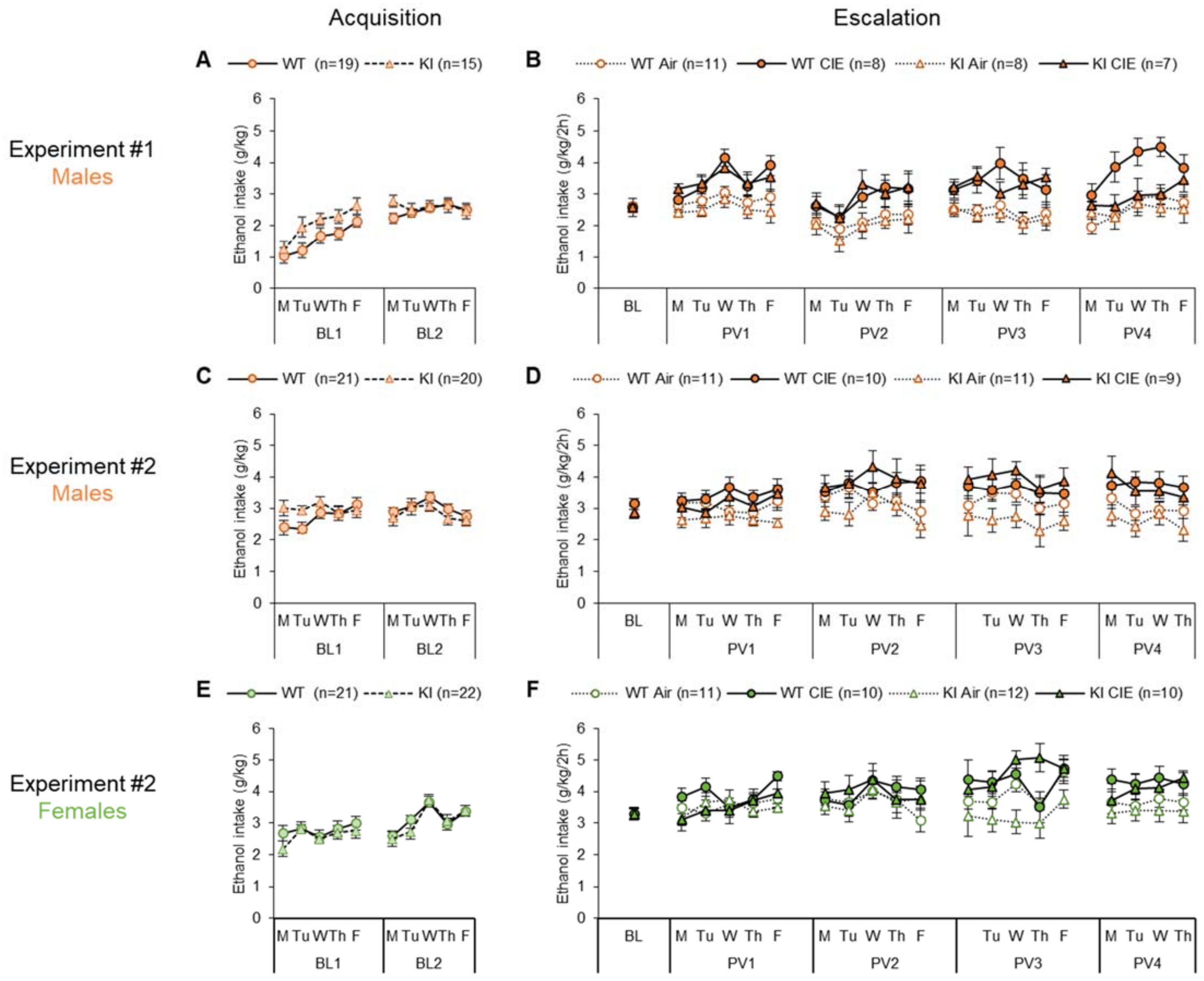


**Supplementary Figure 3. Daily ethanol intake in the CIE-2BC model.** BK α K361N WT and KI mice were given access to voluntary alcohol consumption in 2-h two-bottle choice sessions prior to (Acquisition, **A**, **C**, **E**) and in-between (Escalation, **B**, **D**, **F**) weeks of chronic intermittent ethanol (CIE) vapor inhalation. A first experiment was conducted in males only (**A-B**) and a repeat experiment included both males (**C-D**) and females (**E-F**). Data are shown as mean ± s.e.m. and sample sizes are indicated in the legend of each graph. Statistical analysis was conducted on weekly averages, results are reported in the main text and Figure 4.

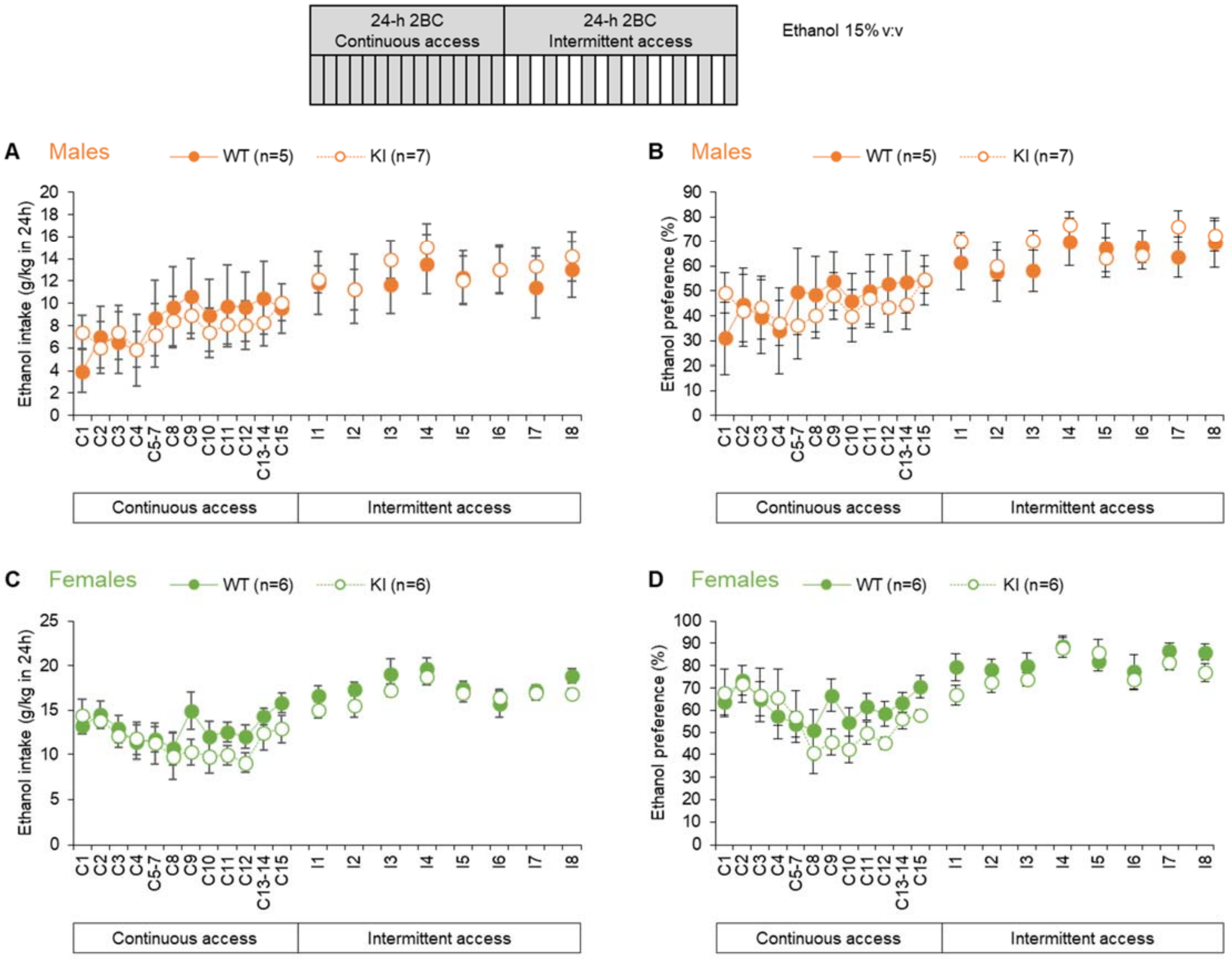


**Supplementary Figure 4. Daily ethanol intake and preference under continuous and intermittent access.** BK α K361N WT and KI males (**A-B**) and females (**C-D**) were given continuous access to voluntary alcohol consumption (two-bottle choice) before switching to an intermittent schedule of access (24 h, three times per week). Ethanol intake (**A**, **C**) and preference (**B**, **D**) are shown for each 24-h period of alcohol access. Data are shown as mean ± s.e.m. and sample sizes are indicated in the legend of each graph. Statistical analysis was conducted on the last 24-h period of each phase, results are reported in the main text and Figure 5.

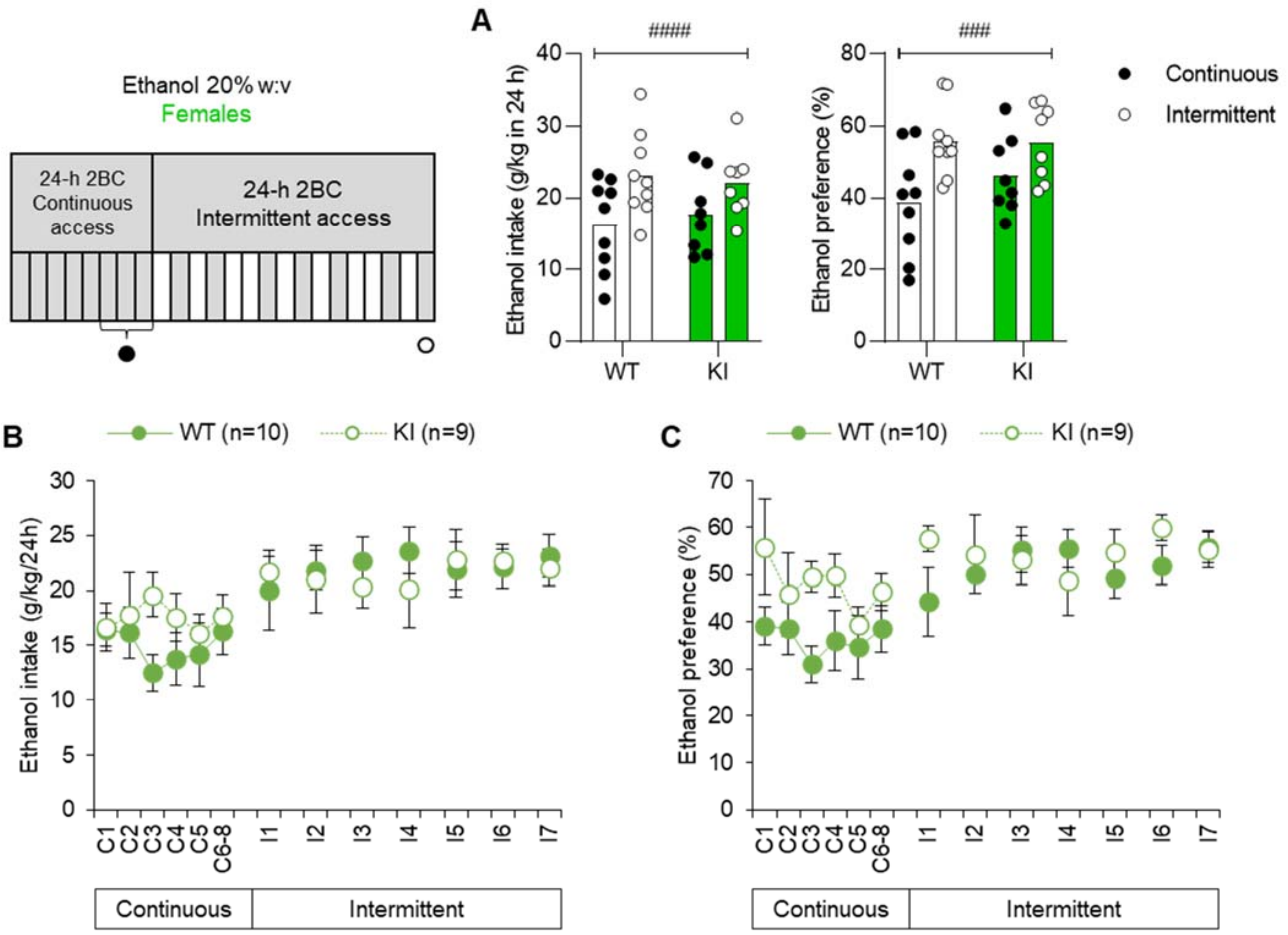


**Supplementary Figure 5. Ethanol intake and preference of wild-type (WT) and BK α K361N knockin (KI) females under continuous and intermittent access to alcohol drinking. A.** Ethanol intake (left) and preference (right) for the last 24-h period of each phase (average of last 72 h for the continuous phase). Main effect of access schedule: $^{###}$, p<0.001. There was no significant effect of genotype on either measure. **B-C.** Ethanol intake (**B**) and preference (**C**) for each 24-h period of alcohol access. Data are shown as mean ± s.e.m. and sample sizes are indicated in the legend.

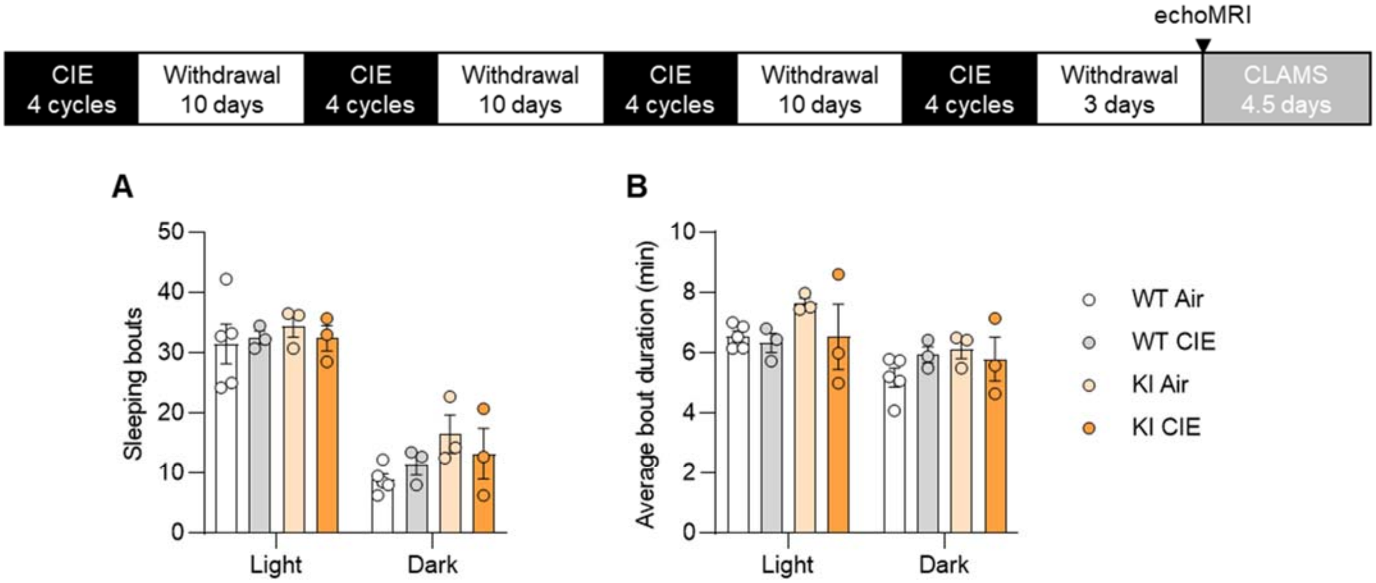


| | Genotype | Vapor | Genotype x Vapor | Phase | Phase x Genotype | Phase x Vapor | Phase x Genotype x Vapor |
|---|---|---|---|---|---|---|---|
| **Sleeping bouts** | $F_{1,11}$=0.02 p=0.90 | $F_{1,11}$=0.4 p=0.54 | $F_{1,11}$=0.02 p=0.90 | $F_{1,11}$=81.1 P<0.0001 | $F_{1,11}$=1.8 p=0.20 | $F_{1,11}$=0.3 p=0.60 | $F_{1,11}$=0.8 p=0.39 |
| **Average sleep bout duration** | $F_{1,11}$<0.01 p=0.97 | $F_{1,11}$=0.2 p=0.66 | $F_{1,11}$<0.01 p=0.94 | $F_{1,11}$=17.9 P=0.0014 | $F_{1,11}$<0.01 p=0.97 | $F_{1,11}$=2.2 p=0.17 | $F_{1,11}$=0.7 p=0.41 |

**Supplementary Figure 6. Actimetry-based sleep measures in wild-type (WT) and BK α K361N knockin (KI) males withdrawn from chronic intermittent ethanol (CIE) vapor inhalation.** Locomotor activity was recorded in metabolic cages 4-8 days after removal from vapor chambers. CLAMS-HC Sleep Detection function used these counts to compute sleep bout numbers (**A**) and duration (**B**). Both measures were higher during the light phase of the circadian cycle, but there was no significant effect of genotype or vapor. Other measures collected in this cohort are presented in Figure 6A-D.

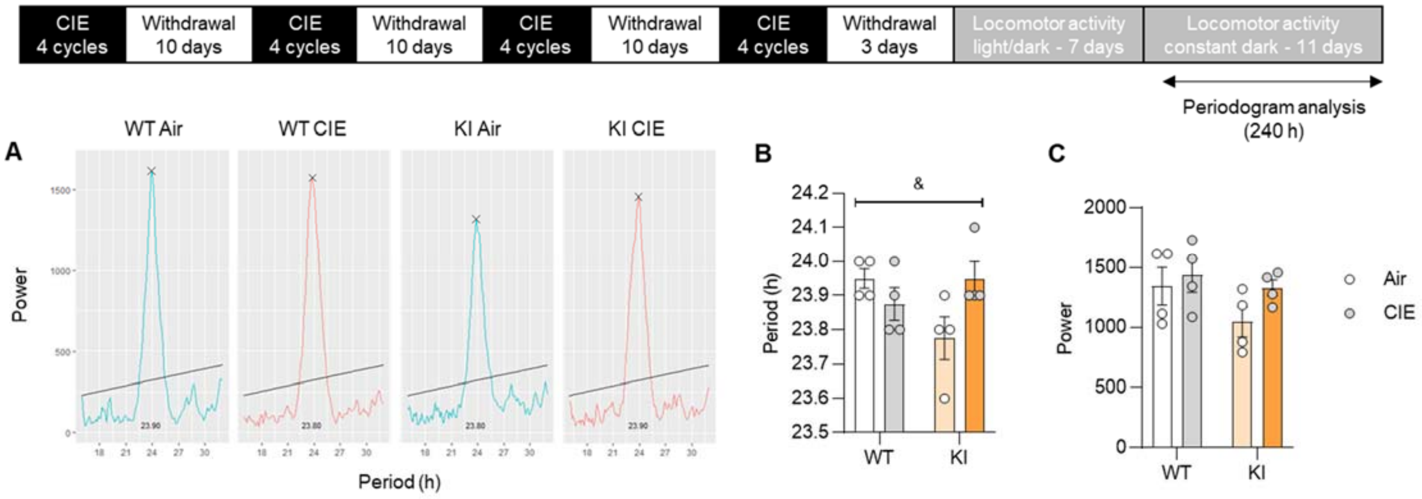


**Supplementary Figure 7. Circadian rhythmicity in wild-type (WT) and BK α K361N knockin (KI) males withdrawn from chronic intermittent ethanol (CIE) vapor inhalation.** BK α K361N WT (n=8) and KI (n=8) mice were exposed to air or chronic intermittent ethanol (CIE) vapor inhalation. Locomotor activity was recorded starting 3 days into withdrawal from CIE (data reported in Figure 6E). After 7 days, mice were switched to constant darkness and periodogram analysis (representative plots shown in **A**) was used to determine the free-running circadian period length (**B**) and relative power (**C**). There was a significant genotype x vapor interaction on the period ($^{\&}$, $p<0.05$) but none of the pairwise comparisons reached significance.
